# Parallel genome-wide CRISPR analysis identifies a role for heterotypic ubiquitin chains in ER-associated degradation

**DOI:** 10.1101/349407

**Authors:** Dara E. Leto, David W. Morgens, Lichao Zhang, Christopher P. Walczak, Joshua E. Elias, Michael C. Bassik, Ron R. Kopito

## Abstract

The ubiquitin proteasome system (UPS) maintains the integrity of the proteome and controls the abundance of key regulators of cellular function by selective protein degradation, but how folding-defective proteins in the secretory system are selected from the large and diverse constellation of membrane and secretory proteins and efficiently delivered to proteasomes in the cytosol is not well understood. To determine the basis of substrate selectivity in human cells, we developed a transcriptional shut off approach to conduct parallel, unbiased, genome-wide CRISPR analysis of structurally and topologically diverse ER-associated degradation (ERAD) clients. Highly quantitative screen metrics allowed precise dissection of entire pathways, enabling identification of unique substrate-specific combinations of recognition and ubiquitin conjugation modules. Our analysis identified cytosolic ubiquitin conjugating machinery that has not been previously linked to ERAD but collaborates with membrane-integrated ubiquitin ligases to conjugate branched or mixed ubiquitin chains to promote efficient and processive substrate degradation.

## Introduction

The ubiquitin (Ub) proteasome system (UPS) maintains the fidelity of the eukaryotic proteome by degrading polypeptides that are unable to correctly fold or assemble into stable native conformations (Amm et al., 2014; Hershko and Ciechanover, 1998). This triage process relies on E3 ubiquitin ligases together with molecular chaperones and substrate adaptors to recognize and tag folding-impaired nascent proteins with polyubiquitin chains, often while still ribosome-associated (Harper and Bennett, 2016; Husnjak and Dikic, 2012; Komander and Rape, 2012). Inefficient triage results in proteinopathies such as amyloidosis and neurodegenerative diseases, while premature degradation of slow-folding but otherwise functional proteins can lead to loss-of-function disorders such as cystic fibrosis (Chen et al., 2015; Guerriero and Brodsky, 2012). How the UPS accurately distinguishes folding-defective proteins from a myriad of nascent, partially folded intermediates in the cytosol is not well-understood; this problem is vastly more complex, however, in compartments that are separated from the cytosolic UPS by bilayer membranes, like mitochondria and endoplasmic reticulum (ER).

Approximately one third of the eukaryotic proteome is synthesized on ribosomes at the cytoplasmic surface of the endoplasmic reticulum (ER) and translocated into or through the lipid bilayer to become membrane or secreted proteins, respectively (Ghaemmaghami et al., 2003). Correct conformational maturation of nascent proteins in the ER is a prerequisite for their export to distal compartments of the secretory pathway (Anelli and Sitia, 2008; Braakman and Hebert, 2013). Proteins that fail to fold or assemble correctly in the ER are degraded by cytoplasmic proteasomes via a process known as ER-associated degradation (ERAD) (McCracken and Brodsky, 1996; Olzmann et al., 2013; Ruggiano et al., 2016). ERAD clients are physically segregated from the UPS and must therefore be partially or completely dislocated across the lipid bilayer to engage the ubiquitin conjugation and proteolysis machinery in the cytosol.

ERAD is organized around membrane-spanning E3 Ub ligases that orchestrate recognition, dislocation and ubiquitin-dependent degradation of substrates by cytoplasmic 26S proteasomes and can be classified into “-L” (lumen), “-M” (membrane) or “-C” (cytosol) based on the orientation of the clients’ folding or assembly lesion or “degron” relative to the ER membrane (**Fig. 1A**). In yeast, two membrane-integrated E3s, Hrd1 and Doa10, handle all ERAD, with Hrd1 mediating ERAD-L and ERAD-M and Doa10 specific for ERAD-C (Carvalho et al., 2006; Denic et al., 2006; Ravid et al., 2006; Vashist and Ng, 2004). By contrast, at least a dozen E3s, including orthologs of Hrd1 (HRD1) and Doa10 (MARCH6), and a large cohort of accessory factors have been linked to ERAD in mammalian cells, reflecting the greatly expanded structural and topological complexity of the secretory and membrane proteomes of metazoans (Christianson and Ye, 2014; Mehnert et al., 2010).

**Figure 1.**
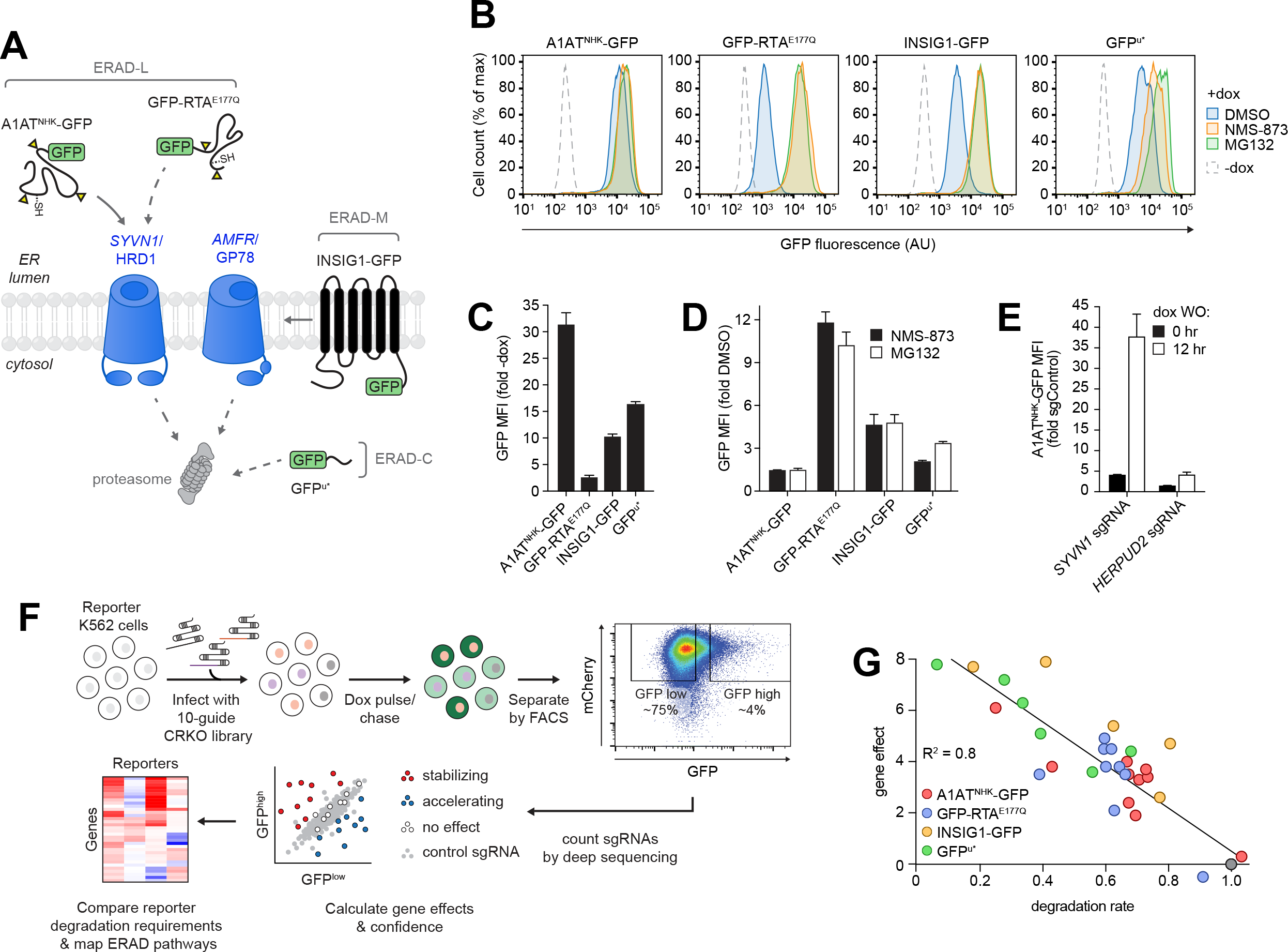
Parallel forward genetic analysis to map substrate-selective ERAD pathways. **(A)** Schematic showing the GFP-tagged ERAD reporters used in this study and the membrane E3 ligases required for their degradation. Yellow triangles indicate sites of N-glycosylation and -SH indicates the location of cysteine residues. A1AT^NHK^-GFP and GFP-RTA^E177Q^ are known (solid arrow) or predicted (dashed arrow) substrates of the HRD1-dependent ERAD-luminal (ERAD-L) pathway; INSIG1-GFP is a substrate of the GP78-dependent ERAD-membrane (ERAD-M) pathway; GFP^u*^ is a predicted substrate of the ERAD-cytosol (ERAD-C) pathway. ERAD-L, -M, and -C clients are targeted to cytosolic proteasomes for destruction. **(B)** ERAD reporter cell lines display increased GFP fluorescence upon UPS impairment. Fluorescence histograms of the indicated K562 ERAD reporter cell lines treated without (dashed line) or with (solid line) doxycycline (dox) for 16 hr before adding DMSO (blue), the p97/VCP inhibitor NMS-873 (orange), or the proteasome inhibitor MG132 (green). Fluorescence was assessed by flow cytometry analysis after 3 hr of inhibitor treatment. **(C)** Steady-state fluorescence reflects reporter half-life. Fold-increase in GFP median fluorescence intensity (MFI) of K562 reporter cells treated with dox for 16 hr. Data are the mean of three independent experiments +/− SEM. **(D)** Effect of inhibitors on steady-state reporter fluorescence. Fold-increase in MFI of dox-induced K562 reporter cell lines treated with 5 μM NMS-873 or 10 μM MG132 for 3 hr. Data are the mean of three independent experiments +/− SEM. **(E)** Doxycycline washout enhances effect of genetic disruption of ERAD machinery. A1AT^NHK^-GFP K562 reporter cells expressing a control sgRNA or a sgRNA targeting the indicated gene were treated with dox for 16 hr. Dox was subsequently washed out (dox WO) and GFP fluorescence was measured by flow cytometry analysis after 12 hr. Data are the mean of three independent experiments +/− SEM. See also Fig. S1. **(F)** Experimental workflow. K562 cells expressing Cas9 and an ERAD reporter were infected with a sgRNA library (10 guides per gene) and the mCherry^+^ infected population was enriched by selecting in puromycin. ERAD reporter expression was induced by treating with dox for 16-20 hr and reporter turnover was induced by removing dox for 0-12 hr. sgRNA-expressing cells were sorted into mCherry^+^/GFP^high^ (brightest ^~^4%) and mCherry^+^/GFP^low^ (dimmest ^~^75%) populations. The frequencies of sgRNAs in each population were determined by deep sequencing and used to calculate an estimated maximal effect (gene effect) and confidence score (gene score) for each gene. Gene effects and scores were compared across screens to identify reporter-specific degradation profiles. **(G)** Gene effects correlate with degradation kinetics. Gene effects measured from sgRNA enrichment in the CRISPR screens were plotted against normalized degradation rates determined by protein synthesis shut off in cells expressing a single sgRNA. Each data point represents a unique gene target identified in one of the four screens. The grey data point indicates cells expressing a control sgRNA. See also Figure S1.

Despite identification of many individual components of the mammalian ERAD machinery, there is little understanding of how they fit together into cohesive modules or networks and, apart from the role of mannose-specific lectins in the recognition of a subset of ERAD-L substrates (Roth and Zuber, 2017; Xu and Ng, 2015), how clients are sorted to these modules. Moreover, while Ub conjugation is essential for dislocation (Baldridge and Rapoport, 2016; Bays et al., 2001; Jarosch et al., 2002; Rabinovich et al., 2002; Stein et al., 2014; Ye et al., 2001), how different E3s and E2s collaborate with each other and with cytosolic UPS and folding machinery in this process for efficient targeting of often highly aggregation-prone clients to proteasomes and p97/VCP has not been systematically investigated, in large measure because of the daunting substrate diversity in the ER and the complexity of the mammalian UPS. Indeed, only a handful of the ^~^ 40 E2 Ub conjugating enzymes and ^~^600 E3 Ub ligases in the human genome have been mapped to substrate-specific ubiquitylation pathways.

To systematically and comprehensively interrogate substrate-selective ERAD modules in human cells we developed a functional genomics pipeline that combines a powerful kinetic assay for protein turnover with an ultracomplex genome-wide CRISPR library and quantitative phenotype metrics. We show that ERAD clients are directed to ER membrane-embedded E3 modules that function with a remarkably specific set of E2s and accessory proteins. Our data reveal a surprising degree of heterogeneity in the role of N-glycans in ERAD client recognition and a remarkably limited set of genes that are shared among different clients. Finally, we identify an unexpected collaboration between membrane-integrated and cytosolic E3s that promotes attachment of ubiquitin with branched or mixed Ub-Ub linkages that are likely to enhance kinetic coupling between dislocation and degradation. These data highlight the power of unbiased, quantitative, multiplex, genome-wide analysis to dissect the highly complex pathways of cellular protein quality control.

## Results

### Parallel, genome-wide CRISPR/Cas9 analysis of genetic requirements for protein degradation

To map the molecular pathways that underlie substrate-selective ERAD, we used pooled genome-wide CRISPR/Cas9 analysis of GFP fusions to four different ERAD and protein quality control substrates (**Fig. 1A**). Two of these, A1AT^NHK^ and INSIG1, are established ERAD-L and ERAD-M clients of the E3s HRD1 (Christianson et al., 2008) and GP78 (Lee et al., 2006), respectively. A third substrate was ricin A chain (K^b^-RTA^E177Q^) engineered with a murine MHC class I (K^b^) signal sequence for efficient cotranslational translocation into the mammalian ER (Redmann et al., 2011) and a point mutation (E177Q) to attenuate its cytotoxity (Ready et al., 1991). RTA normally enters cells as a covalent heterodimer with ricin B chain via endocytosis and is transported retrograde through the secretory pathway to the ER, whereupon it hijacks ERAD dislocons to gain access to the cytosol (Spooner and Lord, 2012). In yeast, RTA dislocation requires Hrd1 and its substrate adaptor Hrd3 (Li et al., 2010), but the E3 required for RTA dislocation in mammals has not been identified. Our study also included GFP^u*^, a variant of a widely-used reporter of cytosolic UPS activity (Bence et al., 2001) that consists of GFP with a C-terminal 16 amino acid CL1 degron (Gilon et al., 1998) with its single lysine residue mutated to glutamic acid. CL1-containing proteins are clients of ERAD-C in yeast (Metzger et al., 2008; Ravid et al., 2006) and mammalian cells (Stefanovic-Barrett et al., 2018).

Basal fluorescence in clonal K562 cell lines expressing these proteins under the control of a tetracycline-On promoter (Gossen and Bujard, 1992) was indistinguishable from untransfected cells, but increased substantially following induction with doxycycline (dox) (**Figs. 1B** and **C**). XBP1 splicing was unaltered upon induction, indicating that expression of these ERAD clients did not activate the unfolded protein response (UPR) and confirming that reporter expression did not exceed the folding or degradative capacity of the ER (**Fig. S1A**). The use of a conditional promoter to acutely drive expression of reporter constructs minimizes cellular adaptations that can result from chronic constitutive expression of folding-impaired proteins (Neal et al., 2018).

To establish a robust selection strategy for phenotypic enrichment, we assessed reporter degradation kinetics and the impact of inhibitors on steady-state fluorescence. The reporters were degraded with half-lives ranging from ^~^10-90 min (**Fig. S1B**) and were strongly stabilized by acutely inhibiting proteasomes with MG132 or p97/VCP with NMS-873 (Hager et al., 2008; Magnaghi et al., 2013) (**Figs. 1B, 1D, S1C**), consistent with the established roles of these enzymes in ERAD (Jarosch et al., 2002; Rabinovich et al., 2002; Ye et al., 2001). Despite near-complete inhibition of reporter turnover (**Fig. S1C**), the effects of these inhibitors on steady-state fluorescence was far more modest and inversely proportion to the reporters’ half-life (**Figs. 1B** and **D**). While the ^~^11-fold effect on the steady-state fluorescence of the very rapidly (t_t/2_ ^~^10 min) degraded GFP-RTA^E177Q^ provides adequate dynamic range for a genetic screen, the 1.4-fold effect on the longer lived (t_1/2_ ^~^90) A1AT^NHK^-GFP does not. This limited dynamic range of steady-state fluorescence measurement has severely limited the sensitivity of previous genetic screens for protein turnover (Christianson et al., 2011; Zhong et al., 2015). We therefore assessed whether transcriptional shutoff could be used as a surrogate “chase” to enhance the dynamic range for phenotypic selection. To promote RNA decay following transcriptional shutoff, the longer-lived reporters A1AT^NHK^-GFP, INSIG1-GFP and GFP^u*^ were expressed without a downstream polyadenylation signal (Hager et al., 2008). Reporter fluorescence remained stable for ^~^120 min following dox washout, reflecting the latency of mRNA decay, then declined at rates proportional to the protein’s intrinsic half-lives (**Fig. S1D**). Strikingly, we found that knocking out *HRD1*, which codes for a E3 Ub ligase known to be essential for A1AT^NHK^ degradation (Christianson et al., 2008), caused a 35-fold increase in fluorescence following dox washout, compared with only ^~^3.5-fold increase in steady-state fluorescence in the presence of the inducer (**Figs. 1E** and **S1E**). Knockout of *HERPUD2*, a gene of unknown function that is required semi-redundantly with *HERPUD1* for A1AT^NHK^ degradation (Huang et al., 2014; Sugimoto et al., 2017), led to a more modest, ^~^4.5-fold increase following dox washout, compared to only ^~^1.3-fold steady state increase, recapitulating the partial stabilization of ERAD-L clients previously observed upon siRNA-directed knockdown (Huang et al., 2014), and suggesting that reporter fluorescence following transcriptional shut off is proportional to protein half-life. These findings confirm that transcriptional shut off substantially enhances the dynamic range of fluorescence-based screens for degradation of UPS substrates.

To identify genes involved in reporter degradation, we developed a functional genomics pipeline (**Fig. 1F**) using a high complexity, full-genome single-guide RNA (sgRNA) library (10 sgRNAs/gene) and statistical metrics to generate an adjusted log_2_ enrichment value (referred to here as the gene effect), a confidence metric calculated from the log-likelihood ratio (referred to here as the gene score), and P-value for each gene (Morgens et al., 2016; Morgens et al., 2017) (**Table S1** and **S2**). The gene effects, calculated from sgRNA enrichment in the sorted populations, were well-correlated with actual reporter degradation rates measured by standard translational shut off analysis in reporter cells expressing single sgRNAs, establishing that screen metrics accurately and quantitatively reflect experimentally determined protein half-lives across a wide range of reporters and effect sizes (**Fig. 1G**).

### Parallel genome-wide screens reveal exquisite substrate specificity in the UPS

The four screens were highly reproducible (**Fig. S2A**) identifying ^~^400-700 genes for each reporter at a 1% false discovery rate (FDR) (**Fig. S2B**; **Table S1** and **S2**). As expected, all reporters were strongly stabilized by disruption of genes encoding UPS machinery (**Fig. 2A** and **Table S3**), reflecting their intrinsic instability, and by genes required for RNA processing and turnover (**Fig. 2B** and **Table S3**), reflecting a screen design that depends on transcript persistence. Genes involved in mitochondrial function and regulation of gene expression (**Fig. 2B**) are frequently enriched in CRISPR/Cas9 screens using different phenotypic selections and were excluded from further analysis (Adamson et al., 2016; DeJesus et al., 2016; Timms et al., 2016). Genes in pathways not directly involved in ERAD, but required for processes immediately upstream and downstream, including protein synthesis, ER targeting, protein folding, N-glycan biosynthesis, ER-to-Golgi trafficking, and lipid biosynthesis were also identified but, in contrast to the former categories, displayed strong substrate specificity **(Figs. 2A** and **C)**.

**Figure 2.**
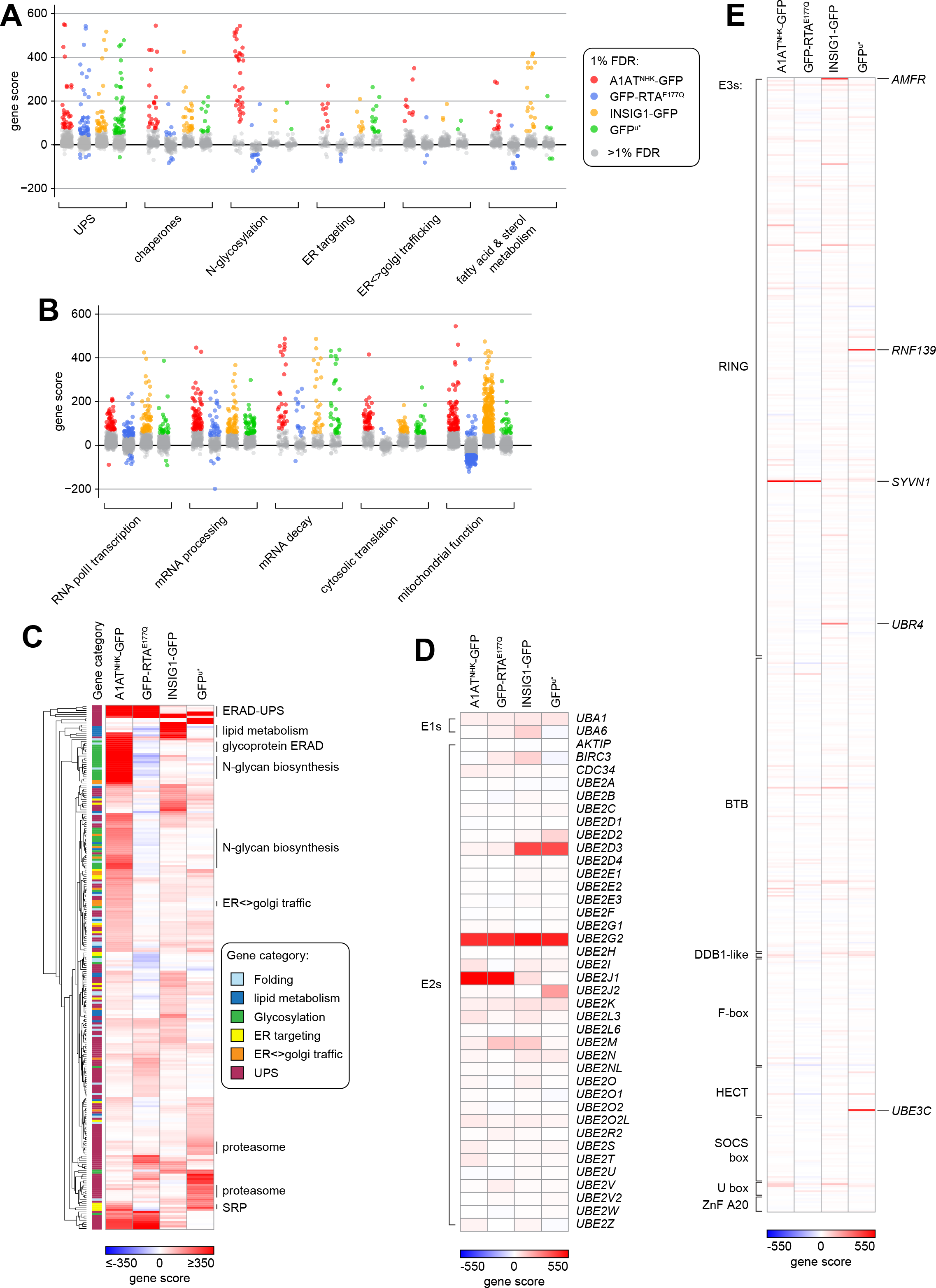
Genome-wide CRISPR analysis identifies genes affecting ERAD reporter stability. (**A-B**) Gene scores for each reporter by functional category. Colored points indicate genes identified at ≤1% FDR and grey points indicate genes at >1% FDR. (A) Includes categories related to ER function and protein quality control; (B) includes categories of genes that likely exert ERAD-independent effects. **(C)** Heat map of genes hierarchically clustered by signed gene scores. A positive score value (red) indicates that the gene was enriched, and a negative value (blue) indicates the gene was disenriched, in the GFP^high^ compared to the GFP^low^ population. Genes in the indicated categories at ≤1% FDR in at least one screen were included in the analysis. **(D-E).** Heat map of signed genes scores for all genes encoding Ub E1 and E2 enzymes (D) and E3s (E). See also Figure S2.

While all four reporters were strongly stabilized by disruption of genes in the UPS category (**Fig. 2A)**, the effects of UPS disruption at the individual gene level were exquisitely specific (**Fig. S2B**), exhibiting distinct substrate-selective “fingerprints”, and only a single UPS gene, *UBE2G2*, was robustly required for degrading all reporters (**Figs. 2C-E**, **S2C** and **D**). Targeting of genes encoding subunits and assembly factors of the 26S proteasome (**Fig. S2C**), and p97/VCP and its cofactors (**Fig. 3A**), also stabilized all four substrates. The lower effect sizes and confidence scores for proteasome and p97/VCP components likely reflects partial loss-of-function phenotypes arising from heterozygous or hypomorphic mutations in these essential genes that survive multiple cell passages before sorting (Estoppey et al., 2017; Parnas et al., 2015; Shi et al., 2015). The remarkable substrate specificity among UPS genes is sharply highlighted by the very limited set of E2s (4 out of 38; **Fig. 2D**) and E3s (5 out of 630; **Fig. 2E**) that were strongly required for substrate degradation. Interestingly, aside from BAP1, which influences reporter expression (data not shown), deubiquitylating enzymes were not found to strongly affect the degradation of any substrate, possibly indicating genetic redundancy or the limited dynamic range of our screen in identifying destabilizing hits (**Fig. S2D**).

**Figure 3.**
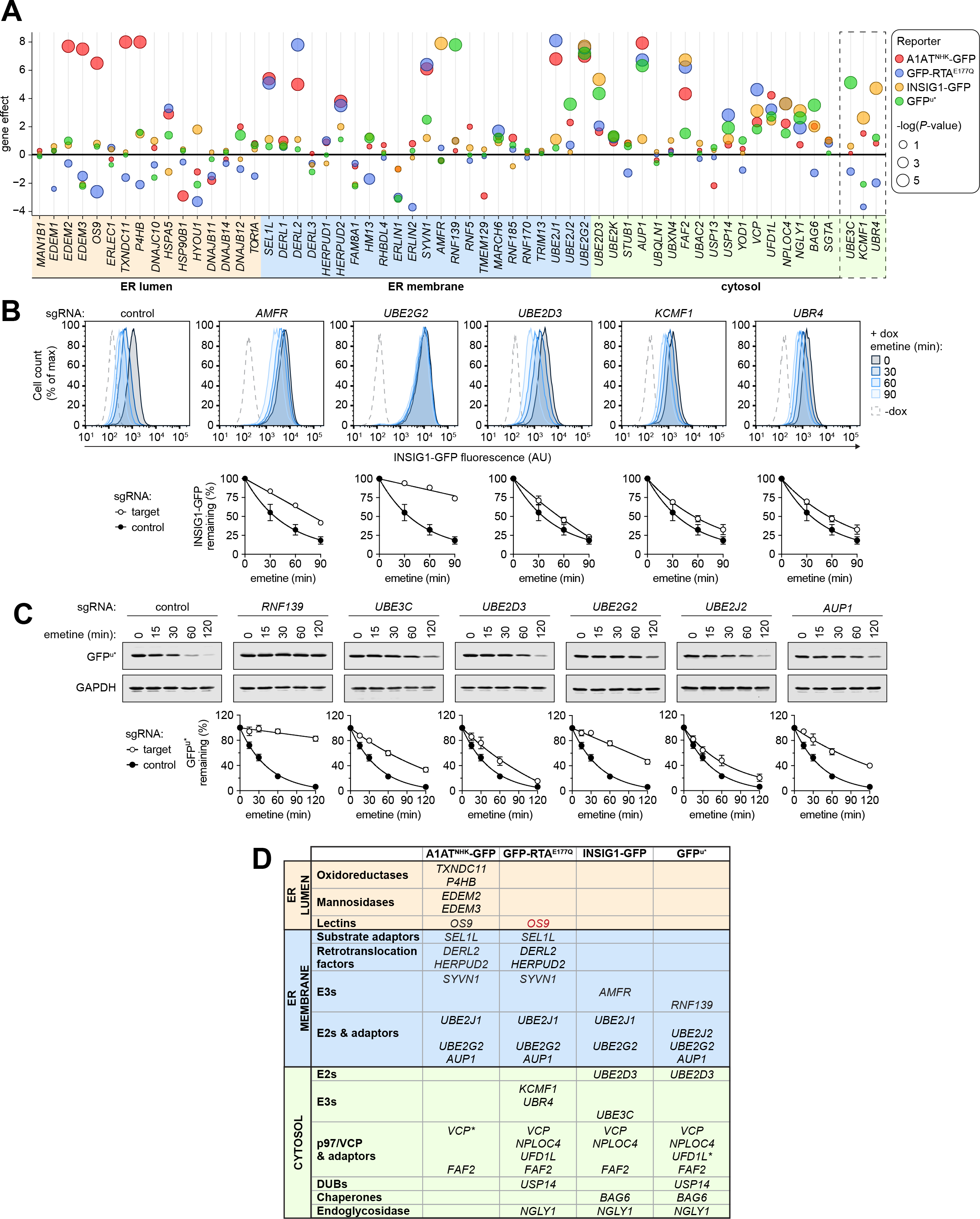
Unique sets of ERAD-related genes are required for degradation of ERAD clients. **(A)** Bubble plot representing the gene effects measured for genes implicated in ERAD. The color of each bubble indicates the reporter screen and the diameter indicates the -log *P*-value. Cytosolic E3 ligases identified in the INSIG1-GFP and GFP^u*^ genetic screens are in the dashed box. **(B)** INSIG1-GFP degradation is impaired by deleting UPS machinery identified in genome-wide CRISPR analysis. Top: fluorescence histograms of reporter cells expressing the indicated individual sgRNAs analyzed by flow cytometry at the indicated times following addition of emetine to shut off protein synthesis. Bottom: Quantification of INSIG1-GFP degradation. Data are the mean of three independent experiments +/− SEM. **(C)** GFP^u*^ degradation is impaired by deleting UPS machinery identified in genome-wide CRISPR analysis. Top: Dox-induced GFP^u*^ reporter cells expressing the indicated individual sgRNAs were treated with emetine for the indicated times and analyzed by SDS-PAGE and immunoblotting with an anti-GFP antibody. Bottom: Quantification of GFP^u*^ degradation. Data are the mean of three independent experiments +/− SEM. **(D)** Summary of the ERAD requirements identified for each reporter. Colored boxes denote topological orientation with respect to the ER membrane as indicated in panel (A). Red font indicates genes that were dis-enriched in the GFP^high^ population relative to the GFP^low^ population. See also Figure S3.

### Substrate specific E3 modules in ERAD

Fine-grained comparison of the significant hits for each of the four reporters (**Figs. 2C** and **3A**) revealed both striking commonalities and differences in their dependencies on UPS and ERAD machinery. INSIG1-GFP was strongly stabilized by disrupting *AMFR*, coding for the membrane-embedded E3 ligase GP78, and *UBE2G2*, confirming a role for this E3/E2 pair in degrading this very hydrophobic integral ERAD-M client (Lee et al., 2006). Unlike HRD1 and TRC8 substrates, which require AUP1 to recruit UBE2G2 to the membrane, this adaptor was not a significant hit for INSIG1-GFP reflecting the fact that GP78 binds and directly activates this E2 via its CUE and G2BR regions (Chen et al., 2006; Das et al., 2009; Liu et al., 2014), underscoring the precision of this unbiased pooled CRISPR approach. By contrast, GFP^u*^ was strongly stabilized by knockout of *RNF139*, which codes for the membrane-integrated E3 TRC8, and by disruption of genes encoding ER-associated E2s, *UBE2J2* and *UBE2G2* (**Figs. 3A** and **C**). TRC8 is required to degrade proteins tethered by hydrophobic anchors to the ER membrane, indicating a role for this ligase in ERAD-C (Boname et al., 2014; Chen et al., 2014). While the vast majority of GFP^u*^ was soluble and partitioned with cytosol, a small fraction was associated with membranes (**Fig. S3A**) and became insoluble upon proteasome inhibition (**Fig. S3B**), consistent with the fact that the CL1* degron is predicted to form an amphipathic α-helix that could allow it to partition weakly with membrane phospholipids.

Surprisingly, in addition to these membrane-integrated factors, INSIG1-GFP and GFP^u*^ were stabilized by disrupting genes encoding the cytosolic E3s, UBR4/KCMF1 and UBE3C, respectively, which have not been previously linked to ERAD, and the cytosolic E2, UBE2D3 (**Figs. 2D** and **E**, **Figs. 3A** and **D**), recently linked to TRC8 and MARCH6-dependent ERAD (Stefanovic-Barrett et al., 2018; van de Weijer et al., 2017). In agreement with these genetic phenotypes, we observed that individual knockouts of the membrane and cytosolic UPS components led to increased steady-state levels (**Figs. S3C-E**) and decreased degradation rates (**Figs. 3B** and **C**) of both reporters.

Curiously, the two nonglycosylated substrates, INSIG1-GFP and GFP^u*^, but not the glycoprotein A1AT^NHK^-GFP, were stabilized by disrupting *NGLY1* (**Fig. 3A** and **D**), which encodes cytoplasmic N-glycanase, an enzyme that removes Asn-linked glycans from dislocated glycoproteins, an activity thought to be required for efficient degradation of glycoproteins by the 26S proteasome (Hirsch et al., 2003; Kim et al., 2006). Mutations in *NGLY1* are linked to a rare hereditary disorder characterized by global developmental delay, movement disorder, hypotonia, and the absence of tears (Bosch et al., 2016; Caglayan et al., 2015; Enns et al., 2014; Heeley and Shinawi, 2015). Because NGLY1 is required for activating the transcription factor, NRF1/*NFE2L1*, which regulates proteasome gene expression in response to stress (Lehrbach and Ruvkun, 2016; Radhakrishnan et al., 2010; Tomlin et al., 2017) it is possible that the modest stabilization of INSIG1-GFP and GFP^u*^ upon NGLY1 knockout could reflect reduced proteasome activity, but it remains unclear why the most heavily glycosylated client, A1AT^NHK^-GFP, was unaffected.

Both A1AT^NHK^-GFP and GFP-RTA^E177Q^ were robustly stabilized by knockout of all components of the well-characterized HRD1/SEL1L complex, confirming the selectivity and sensitivity of our genetic approach and establishing RTA as a HRD1-dependent ERAD client in mammals (**Fig. 3A** and **D**). The extremely short half-life and strict dependence of GFP-RTA^E177Q^ stability on HRD1 is incompatible with reports claiming that RTA escapes proteasomal degradation by virtue of its low lysine content (Deeks et al., 2002). Indeed, HA-RTA^E177Q^ with a lysine-less HA tag was also rapidly degraded and strongly stabilized by deletion of *SEL1L* or by pharmacological impairment of the UPS (**Figs. S3F** and **G**), confirming that this toxin is efficiently recognized and targeted for ubiquitin-dependent proteolysis despite its low lysine content. Although A1AT^NHK^-GFP and GFP-RTA^E177Q^ share a strict dependence on the core membrane-integrated HRD1/SEL1L complex and its associated cytosolic components, they differ strikingly in their dependence on N-glycosylation. A1AT^NHK^-GFP degradation was strongly dependent on the mannosidase complex composed of EDEM2/3 and the oxidoreductases TXNDC11 and PDI that generates the mannose-trimmed Asn-linked glycan which marks proteins for destruction (Clerc et al., 2009; Gauss et al., 2011; Molinari et al., 2003; Ninagawa et al., 2014; Oda et al., 2003; Pfeiffer et al., 2016; Timms et al., 2016), and the lectin, OS9, that recognizes this glycan signal and delivers proteins to HRD1/SEL1L (Christianson et al., 2008; Hosokawa et al., 2009; Quan et al., 2008; Satoh et al., 2010), as well as nearly every gene involved in the formation or transfer of N-linked glycans (**Figs. 4A** and **B**). Unexpectedly, A1AT^NHK^-GFP turnover was insensitive to knockout of *UGP2* and *ALG5*, required to make UDP-glucose and to *ALG6, 8, and 10*, which catalyze the sequential addition of terminal glucoses to core N-glycans, demonstrating that this glycoprotein can be efficiently degraded without first interacting with calnexin/calreticulin. GFP-RTA^E177Q^, by contrast, was robustly *destabilized* by knocking out most genes encoding enzymes required for the synthesis, transfer, processing and recognition of N-glycans, suggesting that these two luminal glycoproteins use vastly different strategies to engage HRD1/SEL1L.

**Figure 4.**
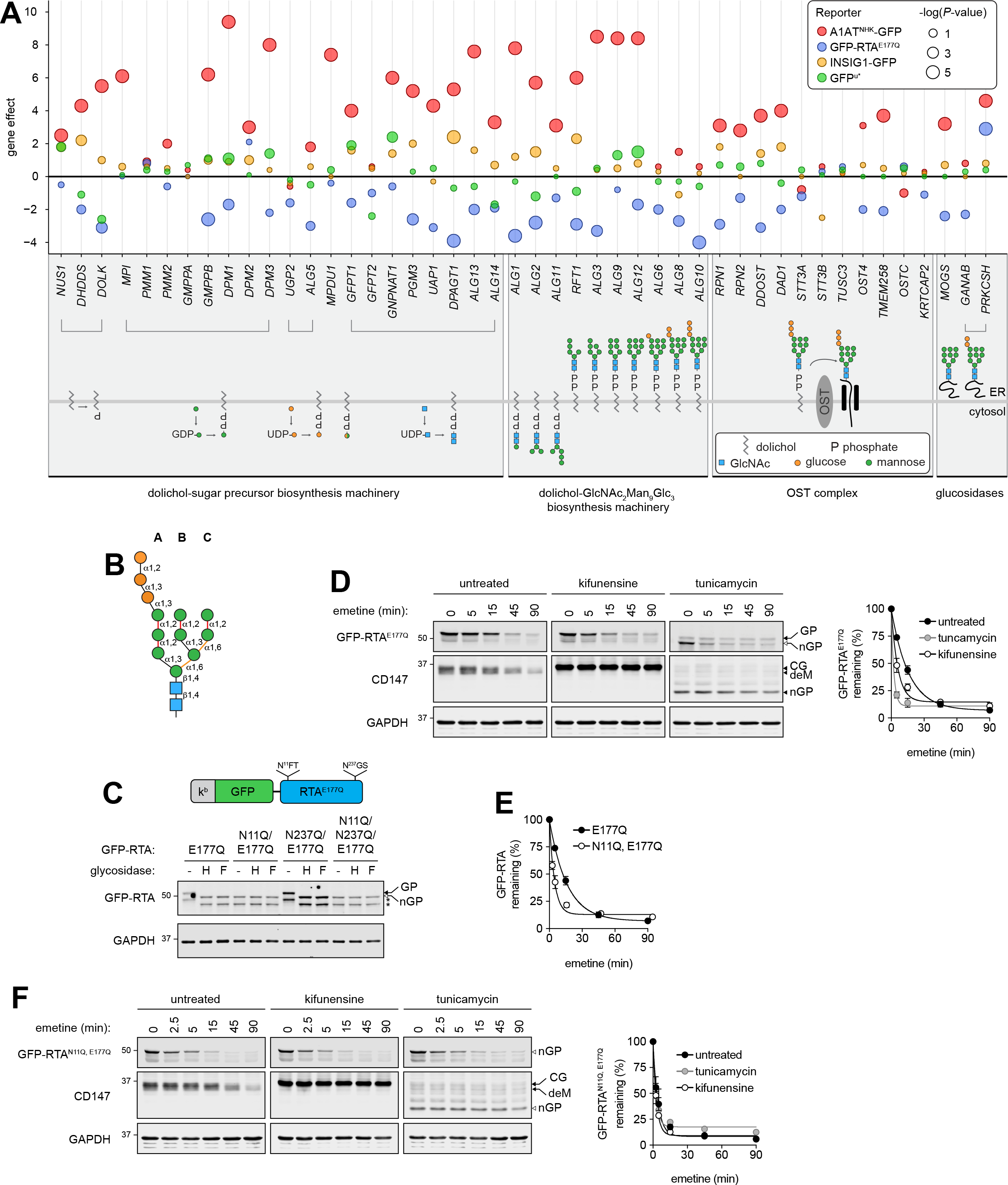
GFP-RTA^E177Q^ is rapidly degraded by evading glycan quality control. **(A)** Bubble plot of gene effects measured for components of the N-glycan biosynthesis pathway. Symbols and colors are as in Figure 3A. **(B)** Structure of the core high-mannose N-glycan. A, B and C-branches and glycosidic linkage topologies are indicated. The α1,2-glycosidic bonds recognized by EDEMs are highlighted in red and the two C-branch α1,6-glycosidic bonds that contact the binding pocket of OS9 are highlighted in yellow. Orange and green circles indicate glucose and mannose residues; blue boxes indicate N-acetyl glucosamine. **(C)** Top: Schematic showing the location and sequence of the two N-glycosylation sequons in GFP-RTA^E177Q^. Bottom: Effect of endoglycosidases on GFP-RTA^E177Q^. Lysates from K562 cells expressing the indicated GFP-RTA^E177Q^ mutants were treated with endo H or PNGase F and analyzed by SDS-PAGE and immunoblotting. Glycosylated (GP) and nonglycosylated (nGP) GFP-RTA^E177Q^ are indicated by filled and open arrows, respectively. Proteolytic cleavage products are indicated by asterisks. **(D)** Left: GFP-RTA^E177Q^ turnover is accelerated by pharmacological inhibition of N-glycosylation or mannosidase activity. Dox-induced GFP-RTA^E177Q^ reporter K562 cells were treated for 6 hr with 2.5 μg/mL kifunensine or for 4.5 hr with 10 μg/mL tunicamycin before inhibiting protein synthesis with emetine. Cells were collected at the indicated times and GFP-RTA^E177Q^ or CD147 turnover was assessed by SDS-PAGE and immunoblotting with an anti-RTA or anti-CD147 antibody. Glycosylated and nonglycosylated forms of GFP-RTA^E177Q^ are indicated as in (C). Arrowheads indicate the core glycosylated (CG), mannose trimmed (deM), and nonglycosylated (nGP) forms of CD147. Right: Quantification of GFP-RTA^E177Q^ turnover. Data are the mean +/− SEM of three independent experiments. **(E)** Nonglycosylated GFP-RTA^E177Q^ is degraded more rapidly than its glycosylated counterpart. Quantification of GFP-RTA^E177Q^ and GFP-RTA^N11Q^’ ^E177Q^ turnover from untreated conditions are the same as in (D) and (F). **(F)** Nonglycolsated GFP-RTA^E177Q^ degradation rate is unaffected by pharmacological inhibition of N-glycosylation or mannosidase activity. Cells expressing dox-inducible GFP-RTA^N11Q^’ ^E177Q^ were treated and analyzed as in (D). See also Figure S4.

### Ricin A chain evades glycan-dependent quality control surveillance

The finding that GFP-RTA^E177Q^ is destabilized by disrupting genes required to generate and recognize the N-glycan “triage” signal was surprising, as GFP-RTA^E177Q^ is predicted to be a glycoprotein and is stoichiometrically glycosylated with a single N-glycan at Asn^11^ (**Fig. 4C**). Individual sgRNAs targeting steps in N-glycan biosynthesis and recognition decreased steady-state levels of GFP-RTA^E177Q^ and accelerated its turnover in a translational shutoff assay (**Figs. S4A, C, D**), confirming that GFP-RTA^E177Q^ destabilization observed in the CRISPR screen reflects an actual increased rate of degradation. Acute pharmacological disruption of N-glycan biosynthesis or mannose trimming with tunicamycin or kifunensine, respectively, accelerated GFP-RTA^E177Q^ turnover to a similar extent as observed upon genetic disruption, while at the same time stabilizing the endogenous, glycan-dependent ERAD substrate CD147 (Tyler et al., 2012) (**Fig. 4D**). This unexpected inhibitory effect of N-glycosylation on GFP-RTA^E177Q^ degradation could be due to the presence of N-glycans on GFP-RTA^E177Q^ itself or on a component of the ERAD machinery. However, the degradation kinetics of GFP-RTA^N11Q,E177Q^, a variant that cannot be glycosylated (**Fig. 4C**), were significantly faster than the glycosylated variant, with the half-life shortened from ^~^12 minutes to less than 2.5 minutes, fully replicating the effects of genetic and pharmacological abrogation of N-glycosylation and mannose trimming (**Fig. 4E**). Moreover, degradation of GFP-RTA^N11Q, E177Q^ was largely unaffected by pharmacological (**Fig. 4F**) or genetic (**Figs. S4B, C, E**) disruption of N-glycosylation, confirming that the acceleration of GFP-RTA^E177Q^ by glycosylation and trimmed mannose recognition is autonomous to the substrate, not the ERAD machinery. Together, these results demonstrate that a demannosylated N-linked-glycan on GFP-RTA^E177Q^ and recognition of this glycan by OS9, which normally *promote* glycoprotein substrate delivery to the HRD1 complex, substantially *retards* the dislocation of this plant toxin. These findings suggest that RTA has evolved a “fast track” access route to the core HRD1/SEL1L dislocon that bypasses the normal route taken by misfolded glycoproteins.

### Heterotypic ubiquitin chains on ERAD-L and ERAD-M clients

The identification of a genetic requirement for cytoplasmic E2s and E3s in INSIG1-GFP and GFP^u*^ degradation (**Fig. 3**) was unexpected, as Ub chain formation on ERAD clients has been thought to rely exclusively on membrane-integrated E3s such as HRD1, GP78 and TRC8, and on membrane-tethered E2s of the “E2G” and “E2J” families (Klemm et al., 2011; Lenk et al., 2002; Spandl et al., 2011). *UBR4* and *KCMF1* encode a large heterodimeric cytosolic E3 complex (Hong et al., 2015; White et al., 2012) recently shown to assemble mixed or branched K11/K48 or K48/K63 linkages (Ohtake et al., 2018; Yau et al., 2017). Conjugation of non-canonical mixed K11/K48 chains to UPS substrates increases the affinity or avidity for binding to p97/VCP and/or the proteasome and the degradation efficiency of aggregation-prone substrates (Meyer and Rape, 2014).

To determine whether atypical Ub linkages are formed on INSIG1, we performed LC-MS/MS analysis on GFP immunoprecipitates from urea-denatured lysates of INSIG1-GFP expressing cells. Two lysines, K33 and K273, of INSIG1 were modified with KGG isopeptide remnants, indicating that INSIG1-GFP is ubiquitylated within its cytoplasmic domain (**Figs. 5A** and **B, Table S4**). In the absence of ERAD inhibitors, INSIG1-GFP was modified with K48-and K11-linked Ub chains (**Fig. 5C**; **Table S4**). Inhibition of the proteasome or p97/VCP led to the additional detection of K6 and K63 Ub linkages. To independently confirm the presence of K11-linked Ub on INSIG1-GFP and to analyze the architecture of these Ub chains, we used a recently described bispecific antibody engineered to bind with high selectivity to branched or mixed Ub chains containing both K11 and K48 linkages (Yau et al., 2017). Ubiquitin conjugates associated with INSIG1-GFP immunoprecipitates from denaturing lysates of dox-induced cells treated with NMS-873 or MG132 were readily detected by immunoblotting using the linkage-specific K11- and K48-antibodies and the bispecific K11/K48 antibody (**Fig. 5D**) but not with control K11/gD or K48/gD antibodies (**Fig. S5A**), suggesting a role for mixed or branched chains containing K11/K48 linkages in degrading this ERAD-M client (**Fig. 5D**). We confirmed modification of INSIG1-GFP by immunopurifying K48 chains or K11/K48 heterotypic chains from denatured cell lysates treated with NMS-873 followed by immunoblotting with a GFP antibody (**Fig. 5E**). These data, beginning with genetic identification of UBR4 and KCMF1 and validated by the presence of both K48 and K11 linked Ub-Ub remnants in immunoisolated INSIG1-GFP and the immunoreactivity of INSIG1-GFP species with bispecific K48/K11 antibodies supports a role for branched or mixed heterotypic Ub linkages in promoting degradation of this ERAD-M client.

**Figure 5.**
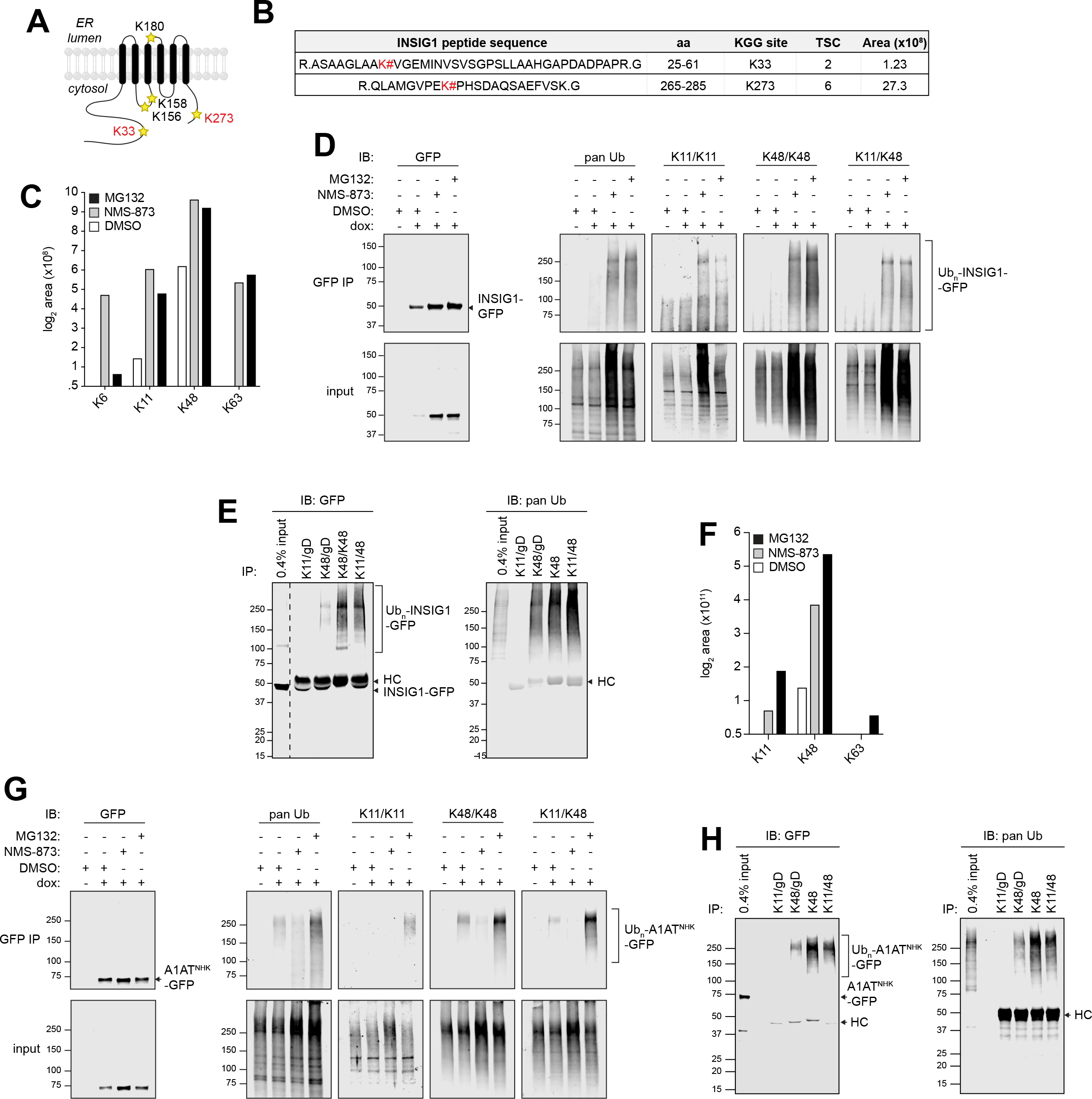
INSIG1-GFP and A1AT^NHK^-GFP are modified with heterotypic K11/K48 polyUb chains. **(A)** Schematic depiction of the location of lysine residues in INSIG1. Lysines modified with Ub are indicated in red. **(B)** INSIG1-GFP ubiquitylation sites identified by LC-MS/MS analysis. INSIG1-GFP was immunoisolated from dox-induced reporter cells treated with 10 μM MG132. # = site of KGG modification; TSC = total spectral counts; Area = integrated MS ion peak intensity. For peptides that were observed more than once, the maximum area measured for that peptide is reported. **(C)** Quantification of INSIG1-GFP-conjugated Ub chains identified by LC-MS/MS. INSIG1-GFP was immunoisolated from cells treated with the indicated inhibitors and Ub KGG peptides were analyzed by LC-MS/MS. **(D)** INSIG1-GFP is modified with K11/K48 mixed or branched Ub chains. INSIG1-GFP was immunoisolated from urea-denatured lysates of dox-induced reporter cells treated with the indicated inhibitors. Ub chain topologies on INSIG1-GFP were detected by immunoblotting with linkage-specific or K11/K48 bispecific antibodies. **(E)** Ub conjugates were immunoisolated using linkage-specific or K11/K48 bi-specific antibodies from denaturing lysates of dox-induced, NMS-873 treated INSIG1-GFP reporter cells. Immunoprecipitated material was separated by SDS-PAGE and INSIG1-GFP was detected by immunoblotting with an anti-GFP antibody. The dashed line indicates removal of irrelevant lanes. HC = IgG heavy chain. **(F)** Quantification of A1AT^NHK^-GFP-conjugated Ub chains identified by LC-MS/MS. A1AT^NHK^-GFP was immunoisolated from cells treated with the indicated inhibitors and Ub KGG peptides were analyzed by LC-MS/MS. **(G).**A1AT^NHK^-GFP is modified with K11/K48 mixed or branched Ub chains. A1AT^NHK^-GFP was immunoisolated from denaturing lysates of dox-induced reporter cells treated with the indicated inhibitors. Ub chain topologies on A1AT^NHK^-GFP were detected by immunoblotting with linkage-specific or K11/K48 bi-specific antibodies. **(H)** Ub conjugates were immunoisolated using linkage-specific or K11/K48 bi-specific antibodies from denaturing lysates of dox-induced, MG132 treated A1AT^NHK^-GFP reporter cells. Immunoprecipitated material was separated by SDS-PAGE and A1AT^NHK^-GFP was detected by immunoblotting with an anti-GFP antibody. HC = IgG heavy chain. See also Figure S5.

The presence of heterotypic, branched/mixed chains on INSIG1-GFP and on GFP^u*^ (see below) prompted us to consider the possibility that HRD1 clients may also be modified with heterotypic Ub chains. Although we did not identify an effect of knocking out cytosolic E3s or E2s on A1AT^NHK^-GFP and GFP-RTA^E177Q^, previous studies showed that the yeast ER-associated E2, Ubc6, together with Doa10, catalyze the formation of K11-linked Ub chains. The mammalian Ubc6 ortholog, UBE2J1, a robust hit for both HRD1 clients in this study, is an integral component of the HRD1/SEL1L complex (Christianson et al., 2011; Hwang et al., 2017; Mueller et al., 2008). While LC-MS/MS analysis of GFP immunoprecipitates from urea-denatured lysates of untreated A1AT^NHK^-GFP K562 cells identified only K48 linked Ub chains, analysis of lysates from NMS-873 or MG132-treated cells revealed the presence of K11 linked Ub chains (**Fig. 5F** and **Table S4**). The presence of K11-K48 branched/mixed chains on the HRD1/SEL1L client, A1AT^NHK^-GFP, was independently confirmed by immunoblotting with linkage-specific K11-and K48-and K11/K48-antibodies (**Figs. 5G** and **S5B**) and by the reciprocal experiment in which K48 and K11/K48 immunoprecipitates were probed with a GFP antibody (**Fig. 5H**). Thus, conjugation of Ub chains with K11 and K48 branched/mixed linkages is a property of two very different ERAD modules, suggesting a broader function in protein quality control than previously anticipated (Komander and Rape, 2012; Yau and Rape, 2016).

### ER and cytosolic UPS machinery conjugates heterotypic ubiquitin chains on GFP^u*^

*UBE3C*, identified in the GFP^u*^ CRISPR screen (**Fig. 3**) codes for a proteasome-associated HECT domain Ub ligase that, like its yeast ortholog Hul5, is thought to function as an “E4”, extending preexisting Ub chains on substrates, an activity proposed to antagonize proteasome-associated DUBs and to promote degradative processivity (Aviram and Kornitzer, 2010; Chu et al., 2013; Crosas et al., 2006; Kohlmann et al., 2008). Impaired GFP^u*^ degradation in *UBE3C*^KO^ cells was rescued by reintroducing wild-type UBE3C, but not a catalytically inactive variant (UBE3C^C1051A^), confirming that E3 ligase activity of UBE3C is required for efficient GFP^u*^ turnover (**Fig. 6A**). Ubiquitylated GFP^u*^ species affinity captured from wild-type cells with Halo-UBQLN1 UBA, an affinity reagent that binds polyubiquitin chains independently of linkage type (Kristariyanto et al., 2015; Ordureau et al., 2014), migrated as a low-mobility smear that further decreased in mobility following exposure to NMS-873 (**Fig. 6B**), consistent with the stabilizing effects of pharmacologic disruption of p97/VCP on GFP^u^ turnover (**Figs. 1B** and **D**). *UBE3C* deletion reduced the relative proportion of high molecular weight (>150 kDa) (Ub)_n_-GFP^u*^ conjugates in the presence or absence of NMS-873 (**Figs. 6B** and **S6A**), consistent with the reported chain extending activity of this enzyme. By contrast, GFP^u*^ was decorated primarily with short chains in wild-type or *UBE3C*^KO^ cells treated with MG132, suggesting that deubiquitylation exceeds chain extension in proteasome-impaired cells and that UBE3C does not contribute to GFP^u*^ oligoubiquitylation. High molecular weight (Ub)_n_-GFP^u*^ conjugates were rescued by re-expressing wild-type UBE3C, but not UBE3C^C1051A^ in *UBE3C*^KO^ cells, confirming that catalytic activity of this E3 is essential for chain extension on oligoubiquitylated GFP^u*^ (**Fig. 6C**).

**Figure 6.**
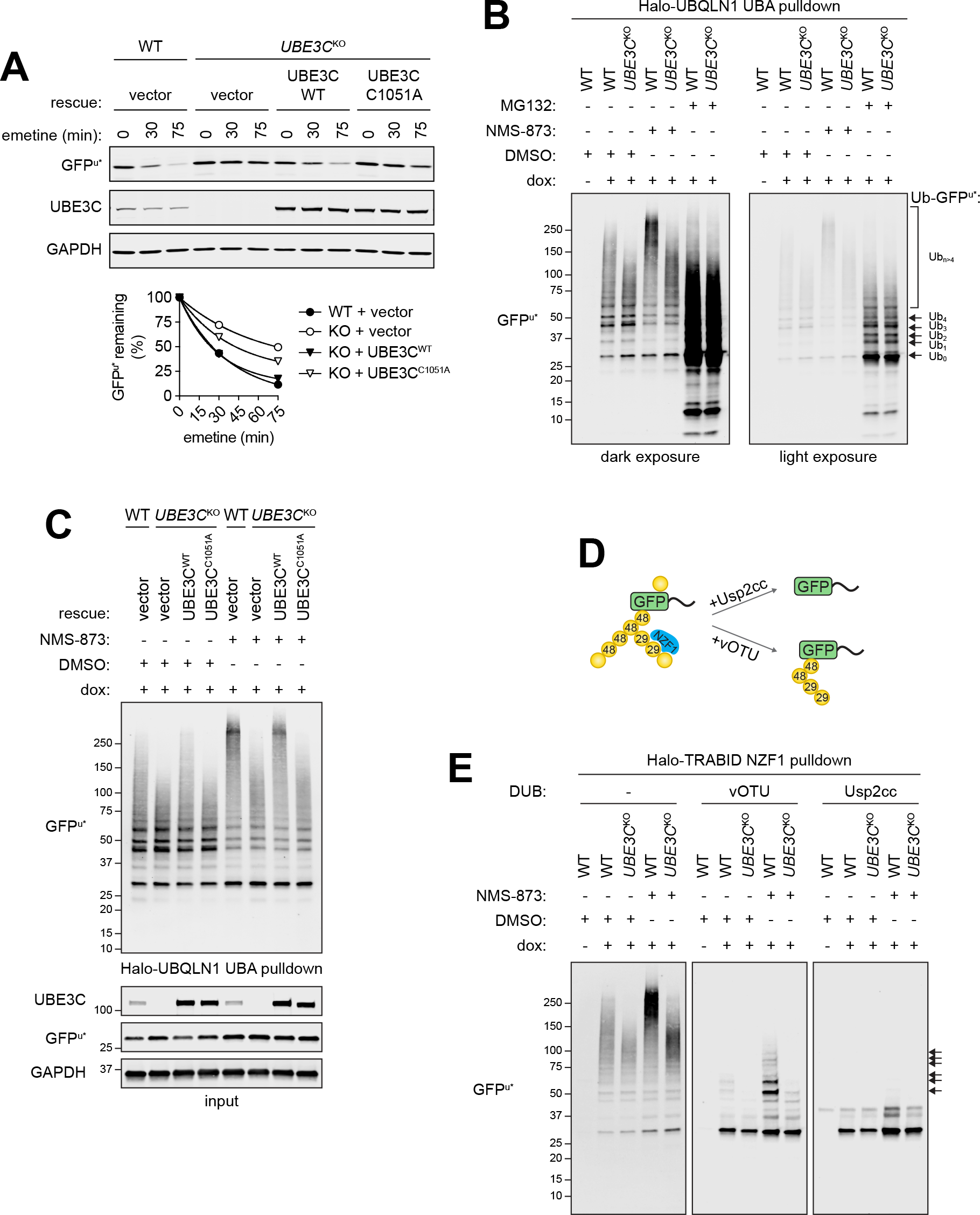
Degradation of GFP^u*^ is promoted by Ub chain diversification. **(A)** GFP^u*^ degradation requires catalytically active UBE3C. Top: Wild type or catalytically inactive mutant UBE3C (UBE3C^C1051A^) was expressed in *UBE3C*^KO^ cells and GFP^u*^ turnover was assessed following inhibition of protein synthesis with emetine for the indicated times. Bottom: Quantification of GFP^u*^ turnover. Data are representative of two independent experiments. **(B)** UBE3C is required for conjugation of high molecular weight Ub conjugates to GFP^u*^. Dox-induced GFP^u*^ reporter cells were treated with the indicated inhibitors for 4 hr. PolyUb conjugates were affinity captured from cell lysates using immobilized Halo-UBQLN1 UBA and analyzed by immunoblotting with an anti-GFP antibody. **(C)** Conjugation of high molecular weight Ub chains to GFP^u*^ requires catalytically active UBE3C. PolyUb conjugates were affinity purified as in (B) from wild type or *UBE3C*^K0^ cells expressing the indicated rescue constructs. Ubiquitylated GFP^u*^ was visualized by SDS-PAGE and immunoblotting with an anti-GFP antibody. **(D)** Schematic diagram for the analysis in (E). Halo-TRABID NZF1 binds selectively to K29-linked Ub chains. **(E)** PolyUb conjugates were affinity captured from GFP^u*^ cell lysates with immobilized Halo-TRABID NZF1 and the presence of K29 Ub linkages on GFP^u*^ was assessed by incubating with the catalytic domain of vOTU, which does not hydrolyze K27 or K29 linkages, or with the core catalytic domain of the nonspecific DUB Usp2 (Usp2cc). Ubiquitylated species were separated by SDS-PAGE and GFP^u*^ was visualized by immunoblotting with an anti-GFP antibody. Arrows indicate polyUb GFP^u*^ species that are resistant to deubiquitylation by vOTU. See also Figure S6.

In addition to catalyzing K48-linked polyubiquitylation, UBE3C can form atypical chains, primarily composed of K29, and to lesser degrees, K11 and K63 linkages (Michel et al., 2015; You and Pickart, 2001). Extension of oligoubiquitin on GFP^u*^ with branched or mixed chains containing K29 linkages could promote its efficient degradation or facilitate degradative processivity by increasing local Ub density, as proposed for K11/K48 branching (Yau and Rape, 2016). To test whether UBE3C makes atypical chains on GFP^u*^, we enriched Ub conjugates from cell lysates by affinity capture with the K29/K33-specific NZF1 Ub binding domain from the DUB TRABID (Kristariyanto et al., 2015; Michel et al., 2015), and immunoblotted for GFP^u*^ (**Figs. 6D** and **E, S6B**). Halo-TRABID NZF1 preferentially captured high MW (Ub)_n_-GFP^u*^ conjugates from control and NMS-873 treated cells (**Fig. 6E**, left panel), suggesting that GFP^u*^ is modified with K29 and/or K33 chains. UBE3C also forms K48 linked chains, which could contribute to the formation of high molecular weight Halo-TRABID NZF1-captured GFP^u*^. To confirm the presence of K29-Ub on GFP^u*^ we treated captured material with the catalytic domain of vOTU, a DUB that cleaves all Ub-Ub linkage types *except* K29 and K27 (Kristariyanto et al., 2015; Ritorto et al., 2014), or the nonspecific DUB, Usp2-cc (Catanzariti et al., 2004; Ryu et al., 2006) (**Figs. 6D** and **6E**, middle and right panels). We observed that GFP^u*^ enriched from wild-type cells contained short polyUb chains that were resistant to vOTU-but not Usp2-catalyzed hydrolysis. The abundance and length of vOTU-protected chains on GFP^u*^ were increased in cells treated with NMS-873. By contrast, Ub conjugates on GFP^u*^ captured from *UBE3C*^KO^ cells treated with or without NMS-873 were efficiently hydrolyzed by both vOTU and Usp2cc. Treatment of Halo-UBQLN1 UBA-captured (Ub)_n_-GFP^u*^ with vOTU and Usp2 produced similar results (**Fig. S6C**). Together, our data reveal that UBE3C catalyzes the extension of oligoubiquitin chains on GFP^u*^ with heterotypic chains containing K29 linkages and promotes efficient proteasome-mediated degradation of a soluble ERAD-C client.

## Discussion

In this paper, we used parallel, genome-wide, CRISPR analysis to elucidate the molecular basis of substrate selective degradation of proteins by ERAD in mammalian cells, allowing us to define three distinct ER membrane-integrated E3 modules that use unique sets of membrane and luminal machinery to efficiently degrade topologically and structurally different ERAD clients with exquisite specificity. Remarkably, this global genomic approach identified a previously unanticipated role for mixed or branched Ub conjugates in the degradation of all three classes of ERAD clients.

### A role for heterotypic ubiquitin chains in ERAD

Degradation of ERAD clients requires coordination of substrate recognition, dislocation, and proteolysis in three cellular compartments. While the mechanisms that direct glycoprotein ERAD-L substrates to HRD1/SEL1L are relatively well understood, far less is known of the mechanisms by which these substrates, once dislocated, are efficiently delivered to 26S proteasomes without aggregating *en route*. Our finding that INSIG1-GFP and GFP^u*^ were stabilized by knocking out membrane-embedded Ub conjugation machinery and cytosolic Ub conjugation machinery, suggest that conjugation of polyubiquitin chains with branched or mixed K11/K48 and K29/K48 linkage topologies contributes to coupling and processivity in ERAD. Our identification of branched/mixed Ub chains on the HRD1 substrate A1AT^NHK^-GFP indicates that such heterotypic chains also likely participate in facilitating HRD1-mediated ERAD, suggesting that conjugation of branched/mixed Ub chains is common to all classes of ERAD substrates and is therefore likely to be a fundamental aspect of ERAD.

Direct Ub conjugation to substrates is essential for dislocation (Baldridge and Rapoport, 2016; Stein et al., 2014), and the positioning of RING domains on ERAD E3s and E2s in a physical complex with dislocons, together with the ability of the E2J class of E2s to attach Ub to serine and threonine residues on ERAD substrates, which are often hydrophobic and deficient in lysines (Shimizu et al., 2010; Wang et al., 2007; Weber et al., 2016), suggests that Ub is conjugated to substrates co-dislocationally (**Fig. 7**). Our findings are consistent with data from yeast mutants suggesting that the ATPase activities of *both* p97/VCP and the 26S proteasome are essential for substrate degradation and that coordination of both activities is necessary to couple dislocation to proteolysis (Lipson et al., 2008; Ye et al., 2001). Although p97/VCP and proteasome engagement are often depicted as discrete, sequential events, we propose that the high local concentration of degradative Ub chains on dislocating ERAD substrates contributes to efficient recruitment of both enzyme complexes, enabling efficient coupling and processivity.

**Figure 7.**
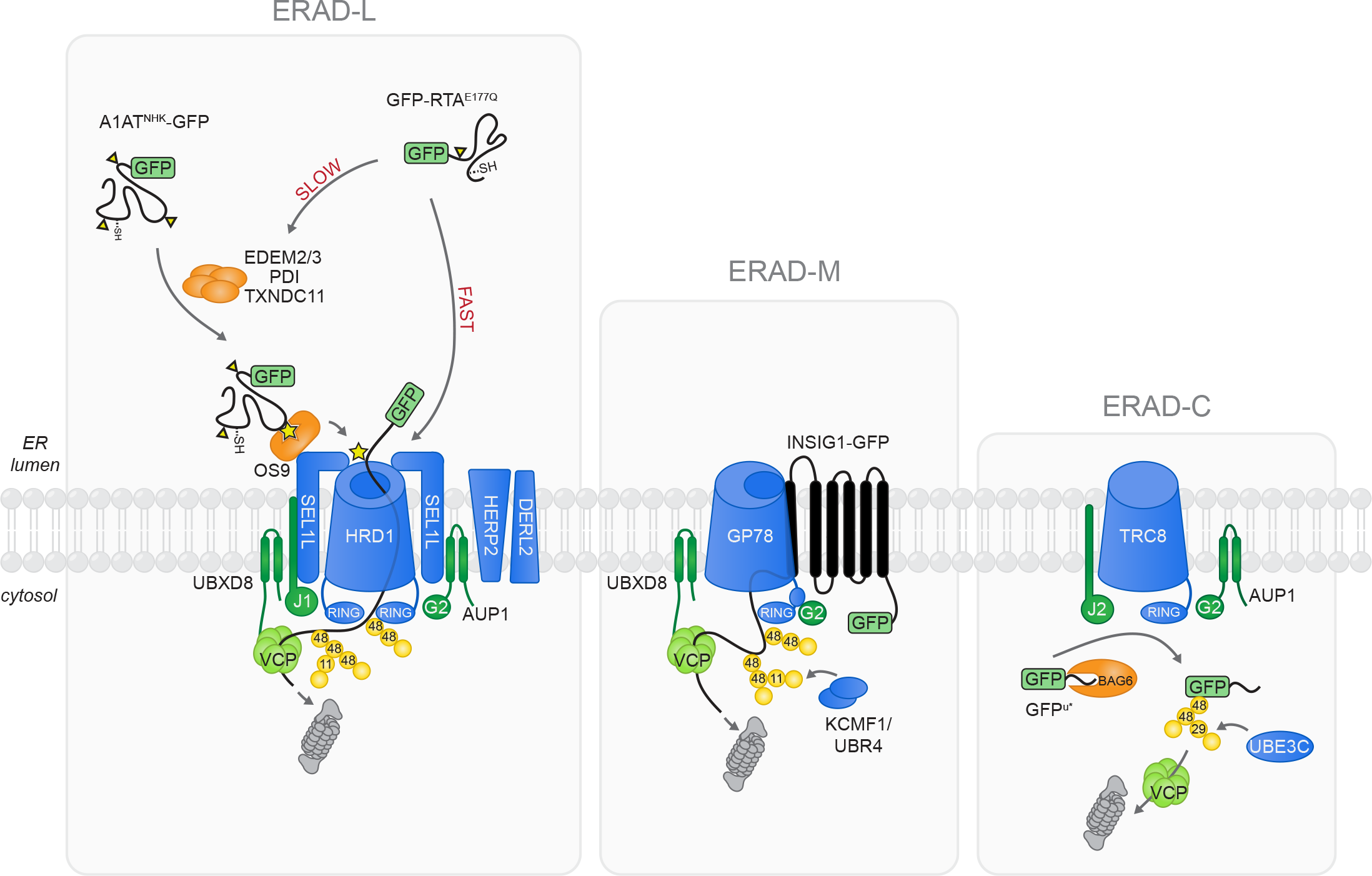
Model of mammalian ERAD-L, -M, and -C pathways mapped through comparative CRISPR/Cas9 genetic screens. (Left) The glycosylated luminal ERAD substrate A1AT^NHK^-GFP requires ER mannosidases and glycan recognition machinery, the membrane-embedded HRD1 dislocation complex, and ER-embedded UPS components for ER exit and delivery to the proteasome, exemplifying the canonical HRD1-dependent ERAD-L degradation pathway. Substrate-specific heterogeneity in the HRD1-dependent pathway is revealed by the luminal glycosylated protein GFP-RTA^E177Q^, which can engage luminal recognition factors for slow delivery to the HRD1 complex or can be rapidly dislocated and degraded by bypassing upstream quality control machinery. Dislocated ERAD substrates are modified with K48 and K11 Ub linkages and targeted for destruction by cytosolic proteasomes. (Middle) The ERAD-M substrate INSIG1-GFP utilizes GP78 for ER dislocation and ubiquitylation. Efficient delivery of this highly hydrophobic substrate to cytosolic degradation machinery may be facilitated by conjugation of K11/K48 chains that increase affinity for p97/VCP and the proteasome. (Right) Efficient degradation of the ERAD-C substrate GFP^u*^ requires conjugation of heterotypic K48/K29 chains via the ER-embedded E3 ligase TRC8 and the cytosolic E3 UBE3C, which may facilitate interactions with proteasome shuttling factors or increase substrate processing at the proteasome.

Heterotypic K11/K48 chains bind to proteasomes and p97/VCP with higher affinity than homotypic chains (Meyer and Rape, 2014) and function as “proteasomal priority signals” that drive efficient degradation of APC/C substrates in mitosis and aggregation-prone mutant substrates of cytoplasmic quality control (Yau et al., 2017). We propose that membrane-embedded E3s collaborate with cytosolic E3s to conjugate these efficient degradation signals to substrates that are either dislocated through or delivered to the ER membrane. The observation that deletion of GP78/*AMFR* and TRC8/*RNF139* resulted in a near complete stabilization of substrates while deletion of *UBR4/KCMF1* or *UBE3C* only partially stabilized GFP^u*^ and INSIG1-GFP, suggests a model in which the membrane-embedded ligases, together with associated UBE2G2 first attach short K48-linked oligoubiquitin chains to substrates followed by extension of these chains by Ub chain diversifying enzymes. The yeast ortholog of UBE3C, Hul5, functions as a Ub chain extender (Crosas et al., 2006) and promotes degradation of aggregation-prone proteins and inefficient proteasome substrates (Aviram and Kornitzer, 2010; Fang et al., 2011; Kohlmann et al., 2008). UBE3C conjugates both K48 and K29 Ub linkages *in vitro* (Kristariyanto et al., 2015; Michel et al., 2015; You and Pickart, 2001). Our data show that UBE3C is also required for the formation of K29 linkages on GFP^u*^ *in vivo.* K29 linkages are predominantly found in mixed or branched heterotypic chains with K48 linkages in cells (Kristariyanto et al., 2015), and while the exact role of K29 linkages is unclear, they accumulate in cells upon proteasome impairment and have been implicated in the ubiquitin fusion degradation (UFD) pathway and ERAD in yeast, suggesting a function in protein degradation (Johnson et al., 1995; Xu et al., 2009a).

UBE3C associates with proteasomes and promotes degradation of a subset of proteins by extending Ub chains and increasing substrate capture; however, whether this is the only function of UBE3C, or whether the chain diversifying function of UBE3C is required, is unclear (Aviram and Kornitzer, 2010; Chu et al., 2013; Crosas et al., 2006; Kohlmann et al., 2008; Lee et al., 2010; Li et al., 2010). A recent study found that in yeast K48/K29 branched chains are added to a UFD substrate downstream of CDC48/p97 and promote binding to the proteasome ubiquitin receptor Rad23p, raising the possibility that UBE3C-dependent chain diversification may facilitate recruitment of proteasomes directly or indirectly (Liu et al., 2017). Future work investigating the order in which types of Ub linkages are added to GFP^u*^, the classes of ERAD substrates modified with K29 linkages, and factors that associate with heterotypic chains, are required to better understand the role of K29 linkages in ERAD.

Genetic identification of the E3 ligase complex composed of UBR4 and KCMF1, implicated in formation of K11/K48 and K48/K63 heterotypic Ub chains (Ohtake et al., 2018; Yau et al., 2017), as promoting turnover of INSIG1-GFP suggested that this ERAD-M substrate may also be modified with atypical Ub linkages. In yeast, deletion of Ubc6 or Doa10 reduces cellular K11 linkages (Xu et al., 2009b) and in mammalian cells depletion of p97/VCP leads to accumulation of ER membrane-associated K11 and K48 linked polyubiquitin conjugates (Locke et al., 2014). However, our study is the first to identify the machinery required to synthesize K11 linkages on ERAD substrates and the first to demonstrate the presence of branched/mixed chains on substrates.

Although we did not identify any other E3s in the CRISPR analysis of A1AT^NHK^-GFP degradation, the presence on this substrate of K11 and K48 linkages suggests that HRD1 itself may be able to conjugate the synthesis of branched Ub chains. We speculate that HRD1, by binding to both UBE2G2 and to UBE2J1, coordinates substrate dislocation and degradation by coupling priming, oligoubiquitylation and extension with branched chains, directly to the engagement of p97/VCP and the 26S proteasome (**Fig. 7**). UBE2J1 (and its yeast ortholog Ubc6) are integral components of the megadalton HRD1 complex, together with UBE2G2 (Ubc7) and its adaptor AUP1 (Cue1) (Christianson et al., 2011; Mueller et al., 2008). A recent cryoEM study (Schoebel et al., 2017) identified the yeast Hrd1/Hrd3 complex as a symmetrical heterotetramer, raising the possibility that it could simultaneously engage two different E2s through its two RING domains, as has been demonstrated for the structurally very different APC/C (Brown et al., 2016). Clearly, much additional structural and biochemical analysis is needed, but the data presented here strongly support the proposition that the three membrane ERAD modules identified here, though significantly different in composition and structure, share a common mechanism that exploits heterotypic Ub chains to facilitate efficient substrate degradation.

### Diverse substrate recognition mechanisms in glycoprotein ERAD

Identification of nearly the entire N-glycan biosynthetic pathway in A1AT^NHK^-GFP degradation illustrates the power of our unbiased pooled CRISPR approach and illuminates how glycans are decoded by ERAD machinery. Most striking is the unexpected finding that GFP-RTA^E177Q^ degradation was strongly *accelerated* by disrupting the same set of glycan biosynthesis, trimming and recognition genes that are essential for degradation of A1AT^NHK^-GFP. Not a typical ERAD substrate, RTA is thought to hijack the ERAD machinery to gain access to the cytosol following endocytic uptake and retrograde migration through the secretory pathway as a disulfide-linked heterodimeric holotoxin (Li et al., 2010; Spooner and Lord, 2012). Identification of genes encoding core HRD1/SEL1L complex components including *HRD1, SEL1L, UBE2G2*, and *UBE2J1*, as protective hits in a growth-based CRISPR/Cas9 screen of ricin holotoxin toxicity (Morgens et al., 2017), strongly supports a role for the HRD1/SEL1L complex in RTA dislocation. However, while both screens identified among the top hits genes required to synthesize core high mannose N-glycans in the ER, the effects of these knockouts in the two screens were completely opposing; disruption of N-glycosylation *accelerated* GFP-RTA^E177Q^ dislocation but strongly *protected* cells from ricin intoxication-most likely because cells deficient in glycan biosynthesis lack the complex glycans with terminal β-linked Gal/GalNAc recognized by ricin’s B-chain and required for efficient endocytic holotoxin capture. Genes required to make these terminal N-glycan modifications in the Golgi apparatus were strong hits for ricin toxicity but were not identified in GFP-RTA^E177Q^ degradation screen. Conversely, plant-produced RTA, unlike the GFP-RTA^E177Q^ construct used here, bears a single complex N-glycan that is not likely to be a substrate for demannosylation by EDEMs or for interaction with OS9, consistent with the observation that the genes encoding these proteins, strongly accelerating for GFP-RTA^E177Q^ degradation, were not captured in the holotoxin screen. The extremely short half-life of GFP-RTA^E177Q^ and strong dependence on ERAD-associated Ub conjugation machinery identified here, together with the robust identification of proteasome subunits as sensitizing hits in the ricin holotoxin screen (Bassik et al., 2013), argue that RTA is efficiently degraded by the UPS, contrasting with the widely held view that the toxin evades degradation by accessing the cytosol independently of ubiquitylation owing to an apparently low lysine content (Deeks et al., 2002; Hazes and Read, 1997; Li et al., 2010). Rather, our data suggest that RTA evolved to maximize its ability to enter the cytosol by exploiting “fast-track” access through the HRD1/SEL1L dislocon that evades the machinery required for quality control surveillance.

### Substrate recognition in ERAD-C

ERAD-C was originally defined in yeast using synthetic transmembrane integrated proteins with folding lesions in the cytoplasmic domain (Carvalho et al., 2006; Ravid et al., 2006; Vashist and Ng, 2004), and many ERAD-C clients are polytopic membrane proteins bearing cytoplasmic mutations (Meacham et al., 2001; Vashist and Ng, 2004). A single E3 in yeast, Doa10 together with Ubc6, Ubc7 and Cue1 ubiquitylate ERAD-C clients (Deng and Hochstrasser, 2006; Ravid et al., 2006; Swanson et al., 2001), but the substrate range for this ligase includes unintegrated tail- or hairpin-anchored membrane proteins (Ruggiano et al., 2016), mis-inserted GPI-anchored proteins (Ast et al., 2014) and soluble cytoplasmic and nuclear proteins bearing, like GFP-CL1, amphipathic C-terminal degrons (Furth et al., 2011; Metzger et al., 2008; Swanson et al., 2001). In metazoans, recognition of this broad range of substrates is diversified among at least three distinct E3s including soluble RNF126, membrane-integrated TRC8 and the Doa10 ortholog, MARCH6 (Boname et al., 2014; Chen et al., 2014; Meacham et al., 2001; Morito et al., 2008; Rodrigo-Brenni et al., 2014; Stefanovic-Barrett et al., 2018; Zelcer et al., 2014). While our study was in progress, a report of a genome-wide insertional mutagenesis screen was published identifying TRC8, AUP1 and UBE2G2 as strong candidates for degrading a GFP^u^ variant, mCherry-CL1 (Stefanovic-Barrett et al., 2018). A fourth hit, MARCH6, was proposed to collaborate with TRC8 to ubiquitylate mCherry-CL1 based on the observation that maximal stabilization of mCherry-CL1, required knockout of both E3 genes (Stefanovic-Barrett et al., 2018). While MARCH6 was also identified in the GFP^u*^ screen it scored below the 1% FDR, while TRC8 knockout lead to nearly complete stabilization.

Cytoplasmic chaperones of the HSP40 and HSP70 families have been reported to contribute to the maintenance of solubility and possibly recruitment of ERAD-C clients to soluble (Meacham et al., 2001) or membrane-associated E3s (Huyer et al., 2004; Metzger et al., 2008). Although our analysis failed to identify cytoplasmic HSP/HSC chaperones, perhaps owing to functional redundancy within this large family, our identification of BAG6 as a hit for GFP^u*^ suggests that this chaperone may contribute to its recruitment to membrane-integrated TRC8. BAG6 is a multifunctional triage factor that plays a central role in sorting proteins with hydrophobic domains either for insertion into membranes or degradation by the UPS. BAG6 binds directly to hydrophobic domains on nascent tail-anchored or type II membrane proteins, promoting their SRP-independent insertion into the ER membrane (Hessa et al., 2011; Leznicki et al., 2010; Mariappan et al., 2010). BAG6 also binds to some ERAD substrates following dislocation, helping to maintain their solubility prior to delivery to the proteasome (Claessen et al., 2014; Wang et al., 2011). Finally, BAG6 can bind to mislocalized SRP-dependent proteins with uncleaved hydrophobic N-terminal signal sequences and, via its Ub like (UBL) domain, facilitate their ubiquitylation by the cytoplasmic E3 RNF126 (Rodrigo-Brenni et al., 2014). Our identification of BAG6 as a stabilizing hit for GFP^u*^ suggests an additional function for BAG6 in recruiting proteins with amphipathic C-termini to TRC8 (**Fig. 7**).

In summary, we report a high precision pipeline to map the genetic pathways by which proteins are degraded by the UPS. The exquisite sensitivity and quantitative readout of this approach allowed us to identify important new players and new roles for known components in ERAD, to define a novel route for RTA intoxication, and to identify ubiquitin ligases that catalyze the formation of branched/mixed atypical Ub linkages on ERAD substrates that are likely to promote processivity. Iterative application of this pipeline to additional substrates can be used to identify the full range of substrate selectivity mechanisms in this complex protein quality control system.

## Materials and Methods

### Cell culture and transfections

K562 human myeloma cells (ATCC) were maintained in RPMI 1640 medium (Corning) supplemented with 2mM L-glutamine (Corning) and 10% FBS (Sigma-Aldrich). Doxycycline (dox)-inducible reporter K562 cell lines were grown in complete RPMI medium supplemented with 200 μg/mL G418 (Thermo Scientific) and 4 μg/mL Blasticidin (Thermo Scientific). HEK293T human embryonic kidney cells (ATCC) were obtained from ATCC and maintained in DMEM (Corning) supplemented with 10% FBS. HEK293 human embryonic kidney cells (ATCC) and HEK293 *SEL1L*^K0^ cell lines (van der Goot et al, In Press, Mol. Cell) were maintained in DMEM supplemented with 10% animal serum complex (Gemini Bio-Products). Cell lines were grown in a humidified incubator at 37°C and 5% CO_2_. All cell lines were routinely tested for mycoplasma infection using a PCR mycoplasma detection kit according to the manufacturer’s instructions (ABM Inc.).

HEK293T cells were transfected using TransIT LT1 (Mirus Bio LLC) and HEK293 cells were transfected using Lipofectamine LTX with Plus Reagent (Thermo Fisher Scientific) according to the manufacturer’s instructions. To transfect K562 cells, 1×10^6^ cells were collected by centrifuging at 1,000 *xg* for 5 minutes. The cell pellet was resuspended in a transfection mix containing 5 μg of plasmid DNA and 100 μL of Opti-MEM medium (Thermo Fisher Scientific) supplemented with 7.25 mM ATP and 11.7 mM MgCl_2_-6H_2_0, moved into a 2 mm electroporation cuvette, and electroporated using a Nucleofector (Lonza) set to program T-016. Transfected cells were cultured for 48-72 hr before being processed for downstream analysis.

### Plasmids

The HA-RTA^E177Q^ expression construct was generated by subcloning kB-HA-RTA^E177D^ (Redmann et al., 2011) (a generous gift from N. Tortorella, Mount Sinai School of Medicine, New York, NY) into pCDNA3.1(+) (Thermo Fisher Scientific) and subsequently mutating residue 177 to a glutamine by site-directed mutagenesis. pBMN2-UBE3C and pBMN2-UBE3C^C1051A^ (Sampson et al., 2013) were generous gifts from T. Wandless (Stanford University, Stanford, CA). pGEX-Halo-TRABID^1-33^ and pGEX6P-vOTU^1-183^ (Kristariyanto et al., 2015) were obtained through the MRC PPU Reagents and Services facility (MRC PPU, College of Life Sciences, University of Dundee, Scotland, mrcppureagents.dundee.ac.uk). The pLVX-Tet-On Advanced vector (Takara Bio) was modified by replacing the CMV promoter with an EF1✓ promoter. The pMCB497-pTRE plasmid used to generate constructs for dox-inducible ERAD reporter expression was made by replacing the EF1 ✓ promoter and cDNA insert from the lentiviral expression plasmid pMCB497 with a custom expression cassette containing pTRE Tight, a multiple cloning site, and pPGK-Blast^R^. The A1AT^NHK^-GFP and GFP^u*^ inserts from previously described constructs (Bence et al., 2001; Christianson et al., 2011) were subcloned into pMCB497-pTRE using a standard restriction enzyme cloning procedure. INSIG1-GFP and GFP-RTA^E177Q^ were generated by replacing the myc tag in pTK-INSIG1-myc (Lee et al., 2006) (a generous gift from J. Ye, University of Texas Southwestern Medical Center, Dallas, Texas) or the HA tag in pCDNA3.1(+)-kB-HA-RTA^E177Q^ with GFP or sfGFP, respectively, using FastCloning (Li et al., 2011). The inserts were subsequently subcloned into pMCB497-pTRE using standard restriction enzyme cloning. GFP-RTA^E177Q^ glycosylation mutants were generated by site-directed mutagenesis of pCDNA3.1(+)-GFP-RTA^E177Q^ and were subsequently subcloned into pMCB497-pTRE. The lentiviral packaging plasmids psPAX2, pMD2.G, pRSV, pMDL, and pVSVG were obtained from Addgene. The SFFV-Cas9-BFP plasmid was described previously (Deans et al., 2016). Individual sgRNA were cloned into the pMCB320 plasmid as previously described (Deans et al., 2016). sgRNA sequences used in this study are listed in Table S5.

### Antibodies and Pharmacological Inhibitors

The primary antibodies used in this study are rabbit α-UBE2G2 mAb (Abcam ab174296; 1:2,500); rabbit α-AUP1 pAb (Proteintech 13726-1-AP, 1:1,000); rabbit α-UBE3C pAb (Bethyl Laboratories A304-122A, 1:1,000); mouse α-UBE2D3 mAb (Abcam ab58251; 1:2,000); mouse α-GFP JL8 mAb (Takara 632381; 1:2,000); rabbit α-GFP mAb (Cell Signaling Technology 2956; 1:1,000); mouse α-PDI mAb (Enzo Life Sciences ADI-SPA-891; 1:1000); rabbit α-VCP pAb (Novus Biologicals NB100-1558; 1:10,000); rabbit α-GAPDH pAb (Millipore ABS16; 1:20,000); mouse α-GAPDH mAb (Proteintech 60004-1-Ig; 1:20,000); rabbit α-alpha tubulin pAb (Abcam ab15246; 1:10,000); mouse α-RTA mAb (BioRad Laboratories MCA2865; 1:2,000); mouse α-CD147 mAb (Santa Cruz Biotechnology Inc. sc-25273; 1:1,000); rabbit α-SEL1L rAb (custom antibody; 1:1,000); rabbit α-Ub pAb (Cell Signaling Technology 3933S; 1:1,000); mouse α-Ub FK2 mAb (Millipore Sigma 04-263; 1:1,000); rabbit α-K48 polyUb, clone Apu2 mAb (Millipore Sigma 05-1307; 1:5,000); human α-K48 polyUb, clone Apu2.07 (Genentech; 0.2 μg/mL); human α-K11 polyUb, clone 2A3/2E6 (Genentech; 0.5 μg/mL); human α-K11/K48 bispecific polyUb (Genentech; 0.2 μg/mL); human α-K11/gD bispecific control (Genentech; 0.2 μg/mL); human α-K48/gD bispecific control (Genentech; 0.2 μg/mL). The secondary antibodies used in this study are goat α-mouse IgG, IRDye 800CW (LI-COR Biosciences 926-32210, 1:10,000); goat α-mouse IgG, IRDye 680LT (LI-COR Biosciences 926-68020, 1:10,000); goat α-rabbit IgG, IRDye 800CW (LI-COR Biosciences 926-32211, 1:10,000); goat α-rabbit IgG, IRDye 680LT (LI-COR Biosciences 926-68021, 1:10,000); goat α-human IgG, IRDye 800CW (LI-COR Biosciences 926-32232; 1:10,000).

Inhibitors used in this study include emetine dihydrochloride hydrate (Sigma-Aldrich), CB-5083 ((Anderson et al., 2015) Cleave Biosciences), NMS-873 ((Magnaghi et al., 2013) Sigma-Aldrich), MG132 (Sigma-Aldrich), tunicamycin (Sigma-Aldrich), kifunensine (Tocris Bioscience), castanospermine (Sigma-Aldrich), E1 inhibitor ((Chen et al., 2011) Millenium Pharmaceuticals, Inc.), and thapsigargin (Sigma-Aldrich).

### Lentivirus packaging, lentivirus infection, and cell line creation

Lentiviral packaging and K562 viral infections were performed as previously described (Deans et al., 2016). Dox-inducible ERAD reporter K562 cells were generated by infecting cells with the modified pLVX-Tet-On Advanced vector. 72 hr after infection, cells were selected in complete medium containing 400 μg/mL G418 and passaged for an additional 10 days before infection with a pMCB497-pTRE vector containing the GFP-tagged ERAD reporter cDNA insert. Infected cells were selected in complete medium containing 7.5 μg/mL Blasticidin and individual clones were isolated by limited dilution cloning.

The clonal cell lines used in this study was screened by immunoblot and flow cytometry analysis and were selected for an absence of ERAD reporter expression when grown in medium lacking dox, homogeneous GFP signal and production of a GFP fusion protein of the expected molecular weight when stimulated with dox, reporter stabilization by the proteasome inhibitor MG132, and normal cell growth rate. Unless otherwise noted, ERAD reporter expression was induced by treating clonal K562 cell lines with dox (Sigma-Aldrich) for 16 hr (0.075 μg/mL for A1AT^NHK^-GFP, 1 μg/mL for GFP-RTA, 0.3 μg/mL for INSIG1-GFP, 0.2 μg/mL for GFP^u*^).

For gene knockout studies, dox-inducible ERAD reporter K562 cell lines were infected with pSFFV-Cas9-BFP lentivirus and cells stably expressing Cas9-BFP were isolated by limited dilution cloning or sorting twice on an Aria II (BD Biosciences) cell sorter equipped with a 405 nm laser. Individual sgRNAs in the pMCB320 vector were introduced into K562 cells by lentiviral infection, cells were passaged for 72 hr after infection, selected in 0.75 μg/mL puromycin (Sigma-Aldrich) for 3-4 days, and recovered in medium lacking puromycin for 3 days. sgRNA-expressing cell lines were assayed within 10 days or were used to generate clonal cell lines by limited dilution cloning.

### Flow cytometry

For all flow cytometry analyses, cells were collected by centrifuging at 1,000 *xg* for 5 min, resuspended in PBS, and placed on ice. At least 20,000 events per sample were analyzed on a LSR II flow cytometer (BD Biosciences) equipped with 405, 488, and 532 lasers or on a FACSCalibur (BD Biosciences) equipped with a 488 laser. Data were analyzed using FlowJo version 10.0.8 (Tree Star).

### Emetine chase

Dox-induced reporter cells were treated with 20 μM emetine (Sigma-Aldrich) for the indicated times and harvested by centrifuging at 1,000 *xg* for 5 min. For flow cytometry analysis, cells were resuspended in PBS and placed on ice. For western blot analysis, cells were washed once with PBS, lysed in 1% SDS lysis buffer (1% SDS, 50 mM Tris-HCl pH 7.5, 150 mM NaCl, 2.5 mM EDTA, 100 mM PMSF, and Roche protease inhibitor cocktail), sonicated for 20 sec on setting 3, and centrifuged at 20,000 *xg* for 15 min. Protein concentration was measured using a BCA Protein Assay Kit (Pierce) and an equal amount of total cell lysate per sample was resolved by SDS-PAGE. Total protein was visualized using REVERT total protein stain according to manufacturer’s instructions (LI-COR Biosciences) and membranes were immunoblotted with the indicated antibodies. Total protein and immunoblots were imaged on an Odyssey CLx imaging system (LI-COR Biosciences). Bands were quantified by densitometry and protein levels at each time point were normalized to total protein or GAPDH. Protein remaining was calculated as a percentage of time 0 and one-phase exponential decay curves were fit using Prism 7 (GraphPad Software).

### Glycosidase treatment

GFP-RTA^E177Q^ reporter K562 cells were lysed in 1% SDS lysis buffer, sonicated for 20 sec on setting 3, and centrifuged at 20,000 *xg* for 15 min. Lysates were heated at 95 °C for 10 minutes and cooled to room temperature before adding Endo H and 10X G5 Reaction Buffer or PNGase F and 10X G7 Reaction Buffer (New England Biolabs, Inc.). Reactions were incubated at 37 °C for 1 hr and then analyzed by SDS-PAGE and immunoblotting.

### Cell fractionation

Triton X-100 (TX-100) soluble and insoluble fractions were isolated from 2×10^6^ dox-induced GFP^u*^ reporter K562 cells by resuspending a PBS-washed cell pellet in 200 μL of 1% TX-100 solubilization buffer (1% TX-100, 50 mM Tris-HCl pH 7.5, 150 mM NaCl, 5 mM EDTA, 20 mM N-ethylmaleimide, 1 mM PMSF, Roche protease inhibitor cocktail). Lysates were rotated at 4 °C for 20 min and then cleared by centrifuging at 20,000 *xg* and 4 °C for 20 min. The TX-100 soluble supernatant was collected and the TX-100 insoluble pellet was washed three times with 300 μL of 1% TX-100 solubilization buffer. The pellet was solubilized by sonicating in 200 μL of 1% SDS solubilization buffer (1% SDS, 50 mM Tris-HCl pH 7.5, 150 mM NaCl, 5 mM EDTA, 20 mM N-ethylmaleimide, 1 mM PMSF, Roche protease inhibitor cocktail) and cleared by centrifuging at 20,000 *xg* and 18 °C for 20 min. Equal volumes of 1% TX-100 soluble and insoluble fractions were analyzed by SDS-PAGE and immunoblotting.

Membrane and cytosolic fractions were isolated from 1×10^6^ dox-induced GFP^u*^ reporter K562 cells by resuspending a PBS-washed cell pellet in 100 μL of 0.04% digitonin buffer (0.04% digitonin, 50 mM HEPES pH 7.5, 150 mM NaCl, 2 mM CaCl_2_, Roche protease inhibitor cocktail). Resuspended cells were incubated at 4 °C for 10 min before centrifuging at 20,000 *xg* and 4 °C for 10 min. The supernatant containing the cytosolic fraction was collected and the pellet was washed three times with PBS before resuspending in 100 μL of 1% TX-100 buffer (1% TX-100, 50 mM HEPES, 150 mM NaCl, Roche protease inhibitor cocktail). The resuspended pellet was incubated at 4 °C for 10 min before centrifuging at 20,000 *xg* and 4 °C for 10 min. The supernatant containing the membrane fraction was collected and the insoluble pellet containing nuclei was discarded. Equal volumes of the cytosolic and membrane fractions were analyzed by SDS-PAGE and immunoblotting.

### Genome-wide CRISPR/Cas9 screen

The 10-sgRNA per gene CRISPR/Cas9 library was synthesized, packaged into lentiviral particles, and infected into ERAD reporter K562 cells stably expressing SFFV-Cas9-BFP essentially as previously described (Morgens et al., 2016). Briefly, each sgRNA sublibrary (9 in total, described previously (Morgens et al., 2017)) and 3rd generation lentiviral packaging plasmids were transfected into 293T cells seeded into one 15-cm tissue culture dish. After 72 hr, lentivirus was harvested and each sublibrary was infected into 35×10^6^ cells. After infection, cells were grown for 72 hr and then selected with 0.75 μg/mL puromycin until the population was ≥90% mCherry^+^. Cells were recovered in medium lacking puromycin for 72 hr, then were collected by centrifuging at 1,000 *xg* for 5 min, resuspended in cell freezing medium (FBS supplemented with 10% DMSO), and stored in aliquots of 5×10^7^ cells.

The ERAD reporter forward genetic screens were performed twice. For genetic screening, cells infected with CRISPR/Cas9 sublibraries were combined to generate two sublibrary pools (sublibrary pool A = Apoptosis and cancer; Drug targets, kinases, and phosphatases; Proteostasis; Unassigned 2; sublibrary pool B = Gene expression; Membrane proteins; Trafficking, mitochondrial, and motility; Unassigned 1; Unassigned 3) and each sublibrary pool was sorted independently. To initiate the screen, a cryopreserved aliquot of each sublibrary was thawed and recovered by passaging in complete RPMI medium for two days and then sublibraries were combined at 1,000-fold coverage. 24 hr later, 350×10^6^ cells were resuspended to a final density of 350,000 cells/mL in complete RPMI medium supplemented with dox. For the GFP-RTA^E177Q^ reporter screen, reporter expression was induced by growing cells in medium containing dox for 20 hr. For the A1AT^NHK^-GFP, INSIG1-GFP, and GFP^u*^ screens reporter expression was induced by growing cells in medium containing dox for 14 hr. Cells were then collected by centrifuging at 1,000 *xg* for 5 min, washed once with warm RPMI medium without supplements, and grown for an additional 12 hr (for A1AT^NHK^-GFP) or 6 hr (for INSIG1-GFP and GFP^u*^) in complete RPMI medium without dox. Cells were harvested by centrifuging at 1000 *xg* for 5 min and then resuspended to a final density of ^~^1.5×10^7^ cells/mL in RPMI medium without phenol red and supplemented with 0.5% FBS, and placed on ice to halt reporter turnover. The cells were separated into a BFP^+^/mCherry^+^/GFP^high^ population containing the brightest ^~^4% of cells and a BFP^+^/mCherry^+^/GFP^low^ population containing the dimmest ^~^75% of cells by sorting on an Aria II equipped with 405, 488, and 532 lasers. For each sublibrary pool, at least 4×10^6^ GFP^high^ and 1×10^8^ GFP^low^ cells were collected.

Genomic DNA was extracted from each cell population using a Qiagen Blood Maxi (for GFP^low^ cells) or Blood Midi (for GFP^high^ cells) kit according to the manufacturer’s instructions. The sgRNAs were PCR amplified from genomic DNA as previously described (Deans et al., 2016) and analyzed by deep sequencing on an Illumina NextSeq. Sequences were aligned to the 10-guide sgRNA library using Bowtie and a likely maximum effect size, score, and *P*-value were calculated for each gene using the casTLE statistical framework (Morgens et al., 2016).

### SDS PAGE and immunoblotting

Proteins were denatured by heating at 65 °C for 10 minutes in 1X Laemmli buffer containing 2% (v/v) 2-mercaptoethanol. Samples were separated by SDS-PAGE and transferred onto Immobilon-FL low fluorescence PVDF (Millipore Sigma) or nitrocellulose membrane (BioRad Laboratories) using a semidry transfer apparatus (BioRad Laboratories). Membranes were blocked with 5% nonfat milk in PBS-T or LI-COR blocking buffer for 30 min −1 hr at room temperature before incubating with primary antibody in 2.5% bovine serum albumin (Sigma-Aldrich) and PBS-T for at least 2 hr. Membranes were washed extensively in PBS-T and then incubated with fluorescence-conjugated secondary antibodies in PBS-T for 45-90 min. Immunoblots were extensively washed in PBS-T and visualized on a LI-COR imager (LI-COR Biosciences) and quantified using ImageStudio Lite v5.0.21 (LI-COR Biosciences).

Ub linkages were detected by immunoblotting with linkage and topology-specific antibodies essentially as described (Newton et al., 2012). Briefly, samples were separated by SDS-PAGE and transferred onto 0.45 μM nitrocellulose membrane by wet transfer at 30 V for 2 hr. Membranes were blocked in 5% nonfat milk in PBS-T for 1 hr before incubating with primary antibody in 5% nonfat milk in PBS-T for 1 hr. Membranes were washed extensively in PBS-T and then incubated with fluorescence-conjugated secondary antibody in 5% nonfat milk in PBS-T for 1 hr, followed by extensive washing in PBS-T and visualizing on a LI-COR imager.

### XBP1 splicing assay

RNA was isolated from 5×10^5^-1×10^6^ cells using an RNeasy mini kit (Qiagen) according to manufacturer’s instructions and cDNA was generated from 500 ng of RNA using SuperScript IV reverse transcriptase (Invitrogen) according to manufacturer’s instructions. XBP1 or PGK1 was amplified from 0.1 μL of cDNA product using the following primers: XBP1 Fw: 5’-TTACGAGAGAAAACTCATGGC-3’; XBP1 Rev: 5’-GGGTCCAAGTTGTCCAGAATGC-3’; PGK Fw: 5’-AAGAACAACCAGATAACAAACAAC-3’; PGK Rev: 5’-GTGGCTCATAAGGACTACCG-3’. PCR products were separated on a 2% agarose/TBE gel and visualized on a UV transilluminator.

### Protein purification

Halo-UBQLN1 4X UBA (Ordureau et al., 2014) recombinant protein was a generous gift from W. Harper (Harvard University, Cambridge, MA). Halo-TRABID NZF1 and vOTU^1-183^ recombinant proteins were prepared as previously described (Kristariyanto et al., 2015). Usp2cc recombinant protein was previously described (Tyler et al., 2012). To conjugate Halo-tagged recombinant proteins to HaloLINK resin, 3.2 mls of HaloLINK slurry (Promega Corporation) was washed three times with binding buffer (50 mM Tris-HCl, pH 7.5, 150 mM NaCl, 0.05% NP-40) before adding 2 mg of recombinant protein. The volume was adjusted to 2 mls by adding binding buffer (for Halo-UBLN1 4X UBA) or binding buffer supplemented with 1 mM DTT (for Halo-TRABID NZF1). Samples were rotated at room temperature for 1 hr or overnight at 4 °C, unconjugated protein was removed by washing the resin five times with binding buffer, and immobilized recombinant proteins were stored at 4C.

### PolyUb affinity capture

Dox-stimulated GFP^u*^ reporter K562 cells were harvested by centrifuging at 1,000 *xg* for 5 min, washed twice with PBS, and lysed in 1% NP-40 lysis buffer (50 mM Tris-HCl pH 7.5, 150 mM NaCl, 1% NP-40, Roche protease inhibitor cocktail) supplemented with 10 mM N-ethylmaleimide. Lysates were incubated at 4°C for 15 min before clearing by centrifuging at 20,000 *xg* at 4°C for 15 min. Protein concentration was measured using a BCA protein assay kit (Pierce), adjusted to 4 mg/mL with 1% NP-40 lysis buffer without N-ethylmaleimide, and then further diluted to 1 mg/mL with Dilution Buffer (50 mM Tris-HCl, pH 7.5, 150 mM NaCl, 5 mM EDTA, Roche protease inhibitor cocktail). For Halo-UBQLN1 UBA affinity captures, 0.5 mg of cell lysate was added to 75 μg of immobilized recombinant protein and rotated at 4°C for 16 hr. Beads were washed three times with High Salt Wash Buffer (50 mM Tris-HCl pH 7.5, 500 mM NaCl, 0.5% NP-40) and once with 10 mM Tris-HCl, pH 7.5. For Halo-TRABID NZF1 affinity captures, 0.5 mg of cell lysate was added to 85 μg of immobilized recombinant protein and rotated at 4°C for 1.5 hr. Beads were washed three times with Low Salt Wash Buffer (50 mM Tris-HCl, pH 7.5, 150 mM NaCl, 0.25% NP-40).

For DUB digestions, affinity captured material was washed 1X with DUB Digestion Buffer (50 mM Tris-HCl, pH 7.5, 150 mM NaCl, 20 mM DTT), liquid was removed, and beads were resuspended in 20 μL of DUB Digestion Buffer and incubated at room temperature for 5 minutes. Beads were subsequently incubated with 2.5 μM vOTU^1-183^ or 3 μg of Usp2cc for 1 hr with gentle shaking at 30°C. Affinity captured material was eluted from the beads by adding Laemmli buffer containing 2% (v/v) 2-mercaptoethanol and incubating at 65°C for 15 min. Eluted proteins and 2% of input were analyzed by SDS-PAGE and immunoblotting.

### Immunoprecipitations

Denaturing immunoprecipitations using linkage-specific antibodies were performed essentially as described (Newton et al., 2012). Briefly, INSIG1-GFP or A1AT^NHK^-GFP reporter K562 cells treated with dox for 16 hr and 5 μM NMS-873 or 10 μM MG132 for 4 hr were resuspended in two cell volumes of Denaturing Lysis Buffer I (8 M urea, 20 mM Tris-HCl, pH 7.5, 135 mM NaCl, 1% TX-100, 10% glycerol, 1.5 mM MgCl_2_, 5 mM EDTA, Roche protease inhibitor cocktail, 2 mM N-ethylmaleimide). Samples were sonicated and rotated at room temperature for 30 min before diluting with an equal volume of Denaturing Lysis Buffer lacking urea. Lysates were cleared by centrifuging at 20,000 *xg* for 15 min and then incubated with Protein A/G Plus Agarose (Pierce) for 1.5 hr at room temperature. 4.5 mg of precleared lysates were incubated with 40 μg of antibody for 12 hr at room temperature and Protein A/G Plus Agarose beads for 1 hr. Beads were washed five times in Urea Wash Buffer (4 M urea, 20 mM Tris-HCl, pH 7.5, 135 mM NaCl, 1% TX-100, 10% glycerol, 1.5 mM MgCl_2_, 1 mM EDTA) and bound proteins were eluted by adding Laemmli buffer containing 2% (v/v) 2-mercaptoethanol and incubating at 65°C for 10 min.

Immunoprecipitations using an α-GFP antibody were performed by lysing in two cell pellet volumes of Denaturing Lysis Buffer II (8 M urea, 20 mM Tris-HCl, pH 7.5, 135 mM NaCl, 1% TX-100, 1.5 mM MgCl_2_, 5 mM EDTA, 50 mM 2-chloroacetamide, 50 μM PR-619, 1 mM PMSF, Roche protease inhibitor cocktail). Lysates were sonicated and rotated at room temperature for 30 min before diluting 1:10 in Denaturing Lysis Buffer II without urea. Lysates were centrifuging at 20,000 xg at 4 °C for 15 min and then incubated with Protein A/G Plus Agarose for 1-2 hr at 4 °C. Pre-cleared lysates were incubated with α-GFP JL8 antibody for 12 hr at 4 °C and Protein A/G Plus Agarose beads for 1.5 hr. Beads were washed four times in Denaturing Lysis Buffer II without urea.

### LC-MS/MS analysis

Immune complexes bound to Protein A/G beads were denatured by heating at 65 °C for 15 min in NuPAGE LDS Sample Buffer (Invitrogen) supplemented with 4.8 mM TCEP (Sigma-Aldrich). 2-chloroacetamide was added to a final concentration of 14 mM and samples were incubated at room temperature in the dark for 1 hr. Samples were separated on a 4-12% Bis-Tris Protein gel (Invitrogen). The gel was fixed and stained in Bio-Safe Coomassie (BioRad Laboratories) and gel slices containing immunoprecipitated proteins were excised and cut into ^~^2mm pieces. Gel pieces were covered with DeDye Solution (50% 25 mM NH_4_HCO_3_/50% Acetonitrile) and incubated with gentle agitation at room temperature for 10 minutes. The de-dying procedure was repeated three times until all coomassie stain was removed and then gel pieces were dried in a Speedvac. Dried gel pieces were covered in 12.5 ng/μL mass spectrometry-grade trypsin (Promega) in 25 mM NH_4_HCO_3_, incubated on ice for 45 minutes, then incubated at 37 °C with gentle agitation for 12 hr. Trypsin-digested peptides were liberated from the gel pieces by incubating with gentle agitation for 25 min in 100-300 μL of Peptide Liberator Solution 1 (25% acetonitrile / 5% formic acid), followed by 25 min in Peptide Liberator Solution 2 (50% acetronitrile / 5% formic acid), followed by 25 min in Peptide Liberator Solution 3 (75% acetonitrile / 5% formic acid). The supernatants containing the liberated peptides were pooled and dried in a Speedvac, then peptides were resuspended in 100 μL of 5% formic acid and desalted with the stage-tip method (Rappsilber et al., 2007).

Desalted peptides were reconstituted in 10 μl of 0.1 % formic acid and analyzed on a LTQ Orbitrap Elite mass spectrometer (Thermo Fisher Scientific, Bremen, Germany) or a Fusion Lumos mass spectrometer (Thermo Fisher Scientific, San Jose, USA). Peptides were separated on a 20 cm reversed phase capillary column (100 μm inner diameter, packed in-house with ReproSil-Pur C18-AQ 3.0 m resin (Dr. Maisch GmbH)). The Orbitrap Elite was equipped with an Eksigent ekspert nanoLC-425 system (Sciex, Framingham, USA) using a two-step linear gradient with 3-25% buffer B (0.2% (v/v) formic acid and 5% DMSO in acetonitrile) for 70 min followed by 25-40% buffer B for 20 min. The Fusion Lumos was equipped with a Dionex Ultimate 3000 LC-system and used a similar two-step linear gradient with 4–25% buffer B (0.1% (v/v) formic acid in acetonitrile) for 80 min followed by 25-45% buffer B for 10 min. INSIG1-GFP samples were analyzed with the Orbitrap Elite system and data acquisition was executed in data dependent mode with full MS scans acquired in the Orbitrap mass analyzer with a resolution of 60,000 and m/z scan range of 340-2,000. The top 20 most abundant ions from MS1 with intensity threshold above 500 counts and charge states 2 and above were selected for fragmentation using collision-induced dissociation (CID) with isolation window of 2 m/z, collision energy of 35%, activation Q of 0.25 and activation time of 5 ms. The CID fragments were analyzed in the ion trap with rapid scan rate. Fragmented ions were dynamically excluded from further selection for a period of 30 seconds. The AGC target was set to 1,000,000 and 5,000 for full FTMS scans and ITMSn scans, respectively. The maximum injection time was set to 250 ms and 100 ms for full FTMS scans and ITMSn scans, respectively. A1AT^NHK^-GFP samples were analyzed with Fusion Lumos system and the data were acquired in top speed data dependent mode with a duty cycle time of 3 s. Full MS scans were acquired in the Orbitrap mass analyzer with a resolution of 120,000 and m/z scan range of 340-1,540. Precursor ions with intensity above 50,000 were selected for fragmentation using higher-energy collisional dissociation (HCD) with quadrupole isolation, isolation window of 1.6 m/z, normalized collision energy of 30%, and precursor ions of charge state +1 excluded. MS2 fragments were analyzed in the Orbitrap mass analyzer with a resolution of 15,000. Fragmented ions were dynamically excluded from further selection for a period of 30 s. The AGC target was set to 400,000 and 500,00 for full FTMS scans and FTMS2 scans. The maximum injection time was set to 50 ms and 200 ms for full FTMS scans and FTMS2 scans.

The resulting spectra were searched against a “target-decoy” sequence database (Elias and Gygi, 2007) consisting of the Uniprot human database (downloaded June 13, 2016) and the sequence of INSIG1-GFP and A1AT^NHK^-GFP, and the corresponding reversed sequences using the SEQUEST algorithm (version 28, revision 12) (Eng et al., 1994) in an in-house software pipeline (Huttlin et al., 2010). The parent mass tolerance was set to +/−10 ppm and the fragment mass tolerance was set to 0.6 Da for Elite data and 0.02 Da for Fusion data. Enzyme specificity was set to trypsin. Oxidation of methionines and GG modification of lysine (114.0429) was set as variable modification and carbamidomethylation of cysteines was set as static modification. Data was filtered to a 1% peptide false discovery rate using a linear discriminator analysis. Precursor peak areas were calculated for protein quantification.

---

**Figure S1.**
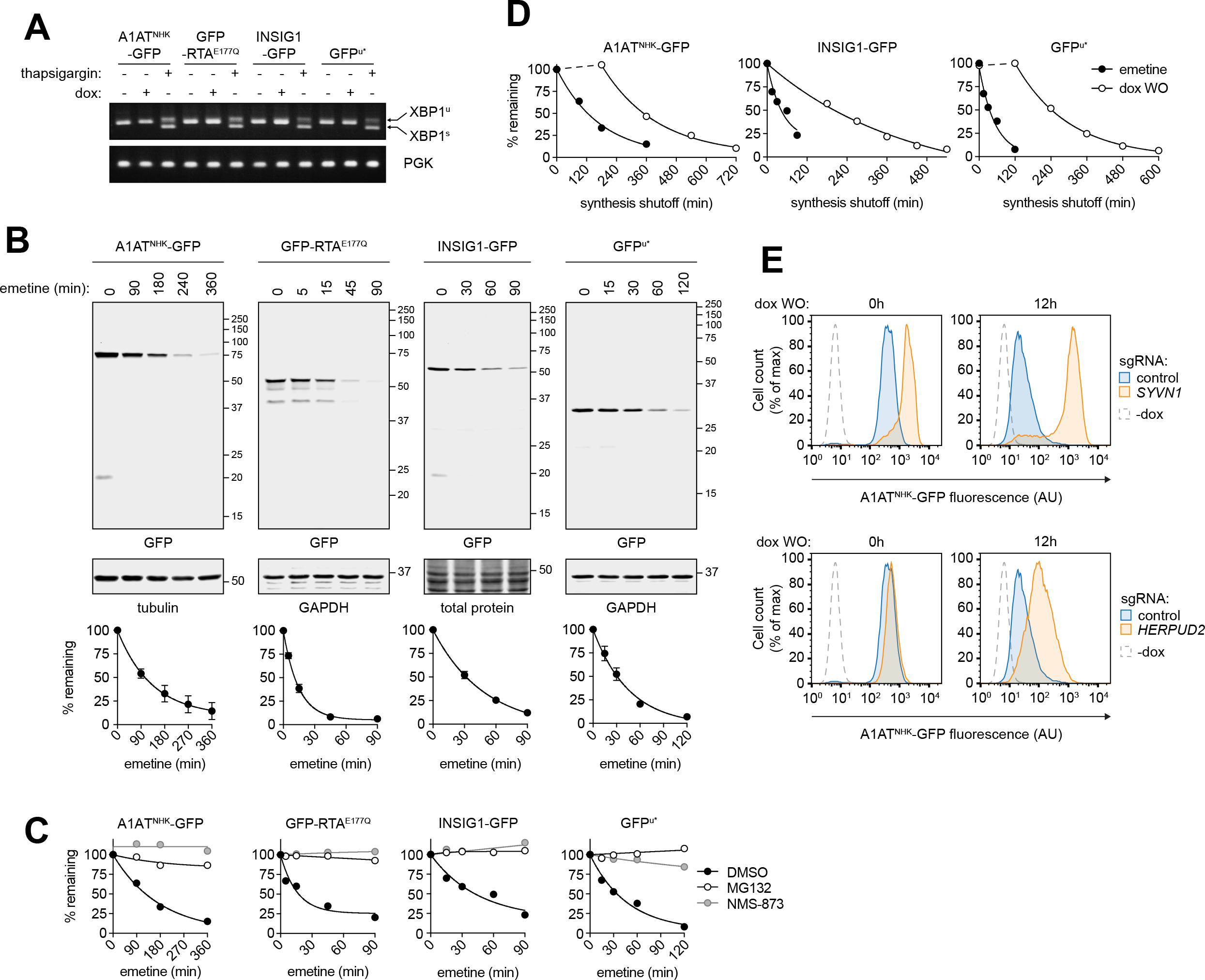
Characterization of ERAD reporter cell lines, Related to Fig. 1. **(A)** Reverse transcription PCR analysis of XBP-1 splicing in ERAD reporter cell lines treated with dox for 16 hr or 1 μM thapsigargin for 1 hr. **(B)** Turnover of GFP-tagged ERAD substrates. Top: Reporter expression was induced with dox and cells were subsequently treated with emetine for the indicated times. Protein was visualized by SDS-PAGE and immunoblotting. Bottom: Quantification of reporter turnover. Data are the mean +/− SEM of three independent experiments. **(C)** Dox-induced reporter cells were treated with DMSO, 5 μM NMS-873, or 10 μM MG132 for 3 hr. GFP median fluorescence intensity (MFI) was measured by flow cytometry at the indicated times after treating with emetine. Data are representative of two independent experiments. **(D)** Reporter decay after translational or transcriptional shutoff. Reporter expression was induced with dox for 16 hr before protein or RNA synthesis was shut off by adding emetine or by removing dox (dox WO), respectively, and GFP MFI was measured at the indicated times by flow cytometry. Data are representative of two independent experiments. **(E)** A1AT^NHK^-GFP K562 reporter cells expressing a control sgRNA (blue), an sgRNA targeting *SYVN1* (orange; top), or an sgRNA targeting *HERPUD2* (orange; bottom) were treated with dox for 16 hr. Dox was subsequently washed out and GFP fluorescence was measured at the indicated times by flow cytometry analysis. See also Fig. 1E.

**Figure S2.**
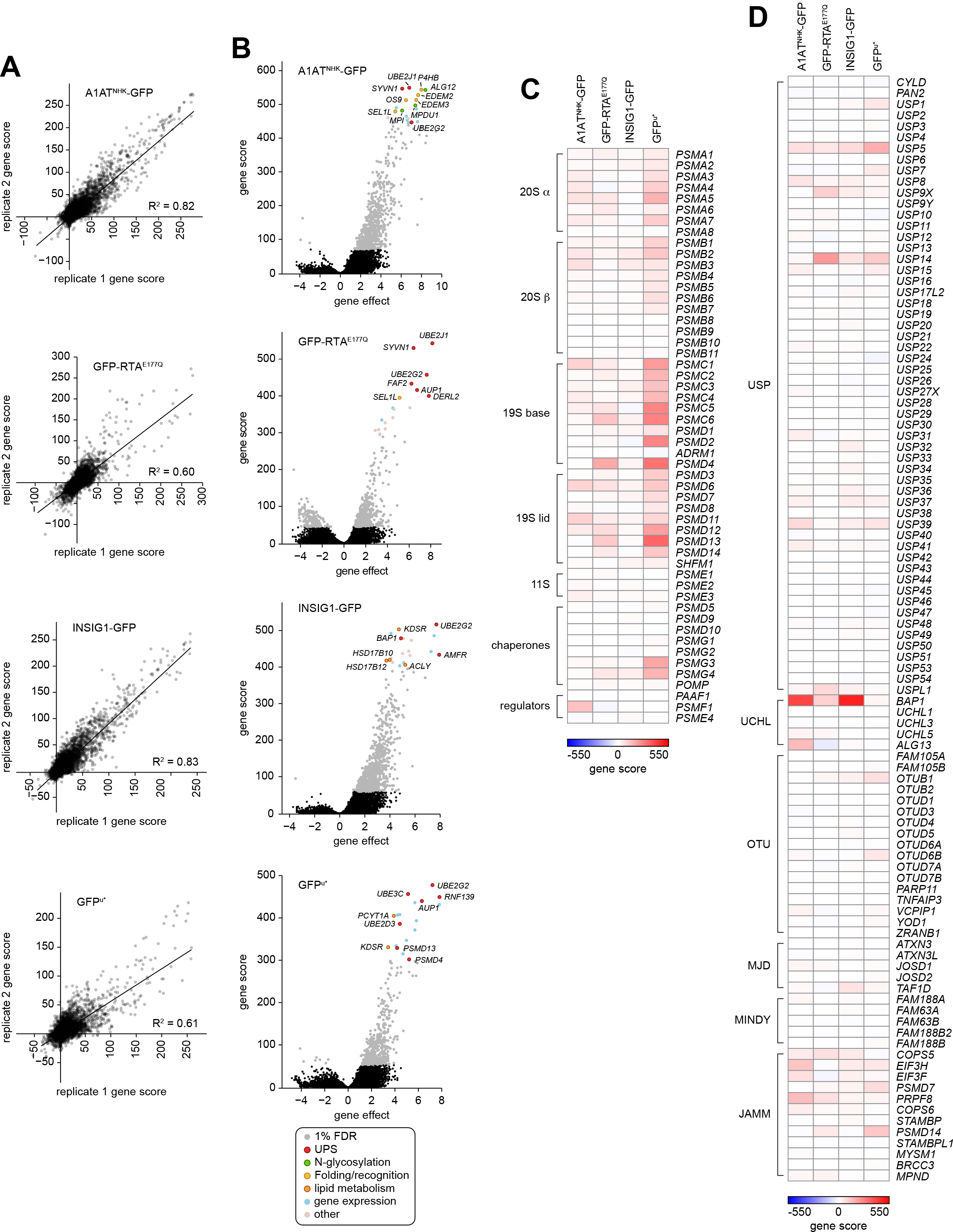
ERAD screens are highly reproducible and identify essential genes, Related to Figure 2. **(A)** Reproducibility of the ERAD genetic screens. Each reporter screen was performed twice and signed gene scores from each replicate were compared. **(B)** Volcano plots of CRISPR analysis for the indicated reporters. Gene effects are plotted against the gene scores and screen hits, identified by applying a 1% false discovery rate (FDR) cutoff, are indicated in grey. The top 20 highest confidence hits from each screen are highlighted. **(C-D).** Heat map of signed genes scores for all genes encoding proteasome subunits, chaperones, and regulators (C) and DUBs (D).

**Figure S3.**
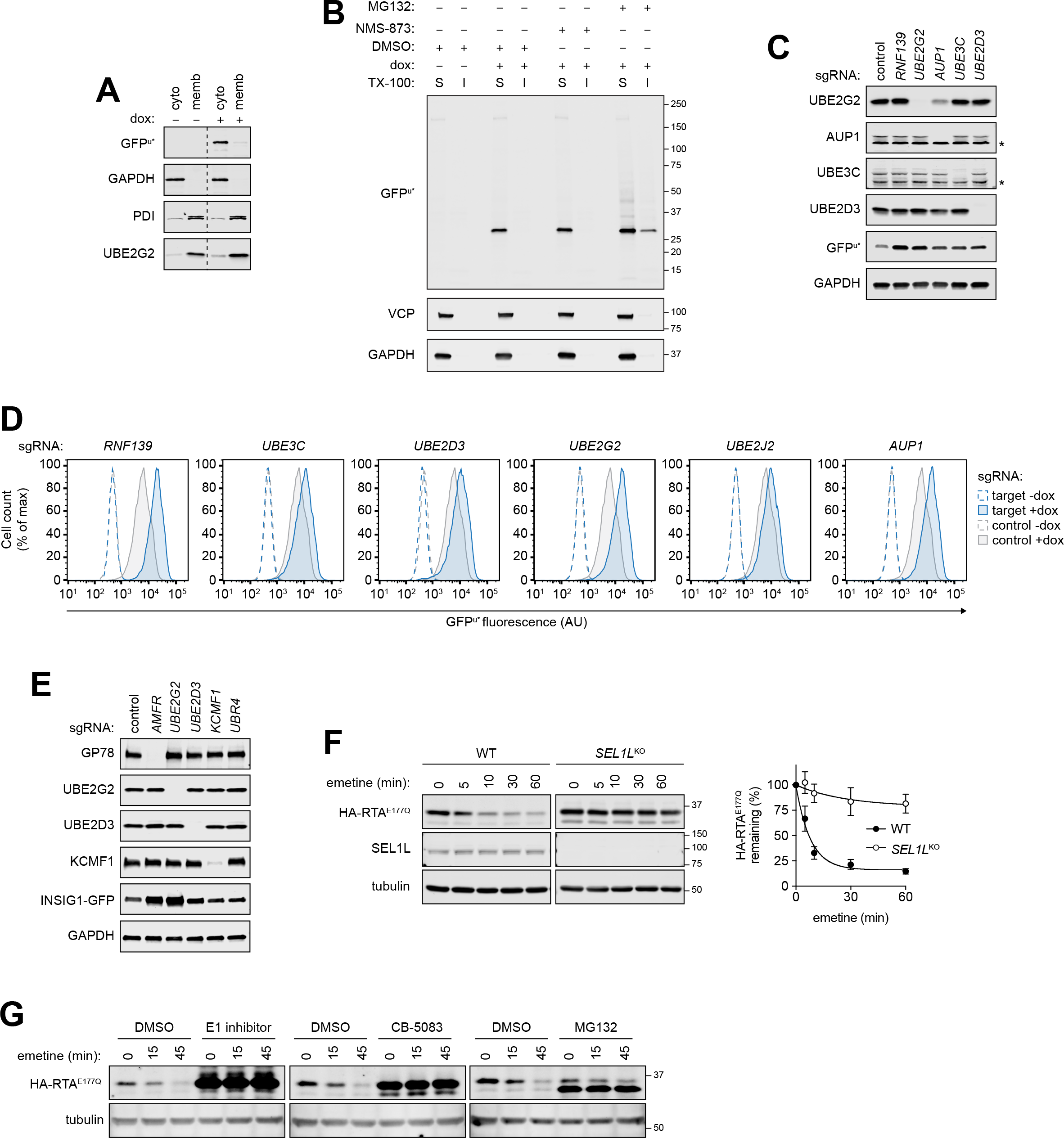
Degradation of GFP^u*^ and RTA^E177Q^, Related to Figure 3. **(A)** GFP^u*^ is cytoplasmic. GFP^u*^ reporter cells were fractionated into cytosolic (cyto) and membrane (memb) fractions and analyzed by SDS-PAGE and immunoblotting with the indicated antibodies. The dashed line indicates removal of irrelevant gel lanes. **(B)** Solubility of GFP^u*^. Control or dox-induced GFP^u*^ reporter cells were treated with the indicated inhibitors for 3 hr. TX-100 soluble (S) and insoluble (I) fractions were analyzed by SDS-PAGE and immunoblotting with an anti-GFP antibody. **(C)** Immunoblot analysis of GFP^u*^ reporter cells expressing sgRNAs targeting the indicated genes. Asterisks indicate nonspecific bands. **(D)** Steady-state levels of dox-induced GFP^u*^ in cells expressing the indicated sgRNAs. GFP fluorescence was determined by flow cytometry analysis. **(E)** Immunoblot analysis of INSIG1-GFP reporter cells expressing sgRNAs targeting the indicated genes. **(F)** Left: Degradation kinetics of HA-RTA^E177Q^. Wild type or *SEL1L*^K0^ HEK293 cells transiently transfected with HA-RTA^E177Q^ were treated with emetine for the indicated times. Reporter turnover was assessed by SDS-PAGE and immunoblotting with an anti-RTA antibody. Right: Quantification of HA-RTA^E177Q^ turnover. Data are the mean +/− SEM of three independent experiments. **(G)** HEK293 cells transiently expressing HA-RTA^E177Q^ were treated with DMSO, 10 μM E1 inhibitor, 5 μM CB-5083, or 10 μM MG132 for 3 hr, followed by emetine for the indicated times. Reporter turnover was assessed by SDS-PAGE and immunoblotting with an anti-RTA antibody.

**Figure S4.**
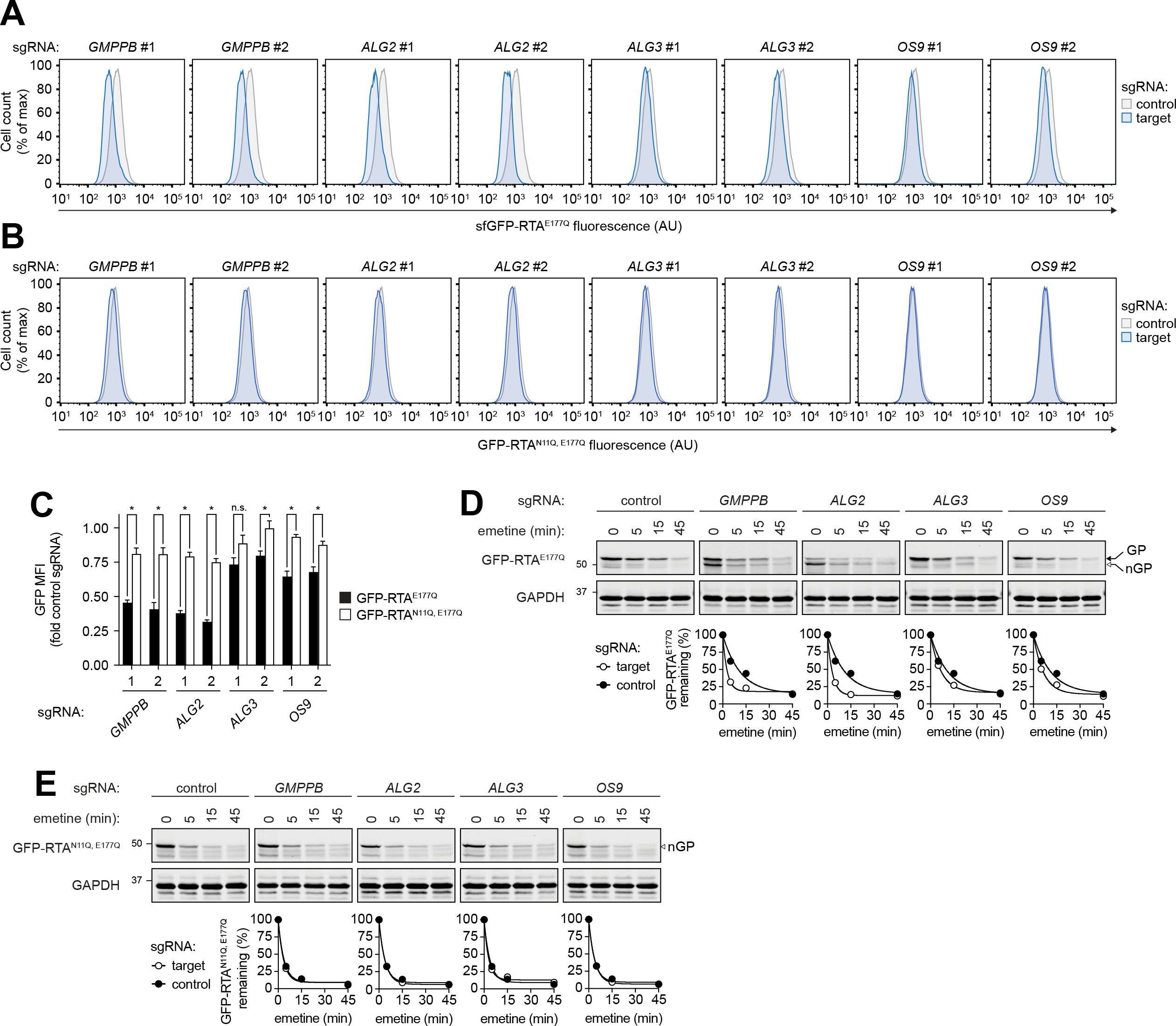
GFP-RTA^E177Q^ degradation is accelerated by deletion of N-glycan biosynthesis and quality control machinery, Related to Figure 4. **(A)** Flow cytometry histograms of steady-state GFP fluorescence in dox-induced GFP-RTA^E177Q^ reporter K562 cells expressing the indicated sgRNAs. **(B)** Flow cytometry histograms of steady-state GFP fluorescence in dox-induced GFP-RTA^N11IQ E177Q^ reporter K562 cells expressing the indicated sgRNAs. **(C)** Steady-state levels of GFP-RTA^E177Q^ and GFP-RTA^N11Q,E177Q^ were measured by flow cytometry analysis in cells expressing the indicated sgRNAs. Bars are MFI +/− SEM from three independent experiments. * indicates *P* ≤ 0.02 (two tailed t-test). **(D)** Top: Dox-induced GFP-RTA^E177Q^ reporter K562 cells expressing the indicated sgRNAs were treated with emetine. Cells were collected at the indicated times and GFP-RTA^E177Q^ turnover was assessed by SDS-PAGE and immunoblotting with an anti-RTA antibody. Glycosylated (GP) and nonglycosylated (nGP) GFP-RTA^E177Q^ are indicated by filled and open arrows, respectively. Bottom: Quantification of GFP-RTA^E177Q^ turnover. Experiment is representative of two independent replicates. **(E)** Cells expressing dox-inducible GFP-RTA^N11IQ E177Q^ and the indicated sgRNAs were treated and analyzed as in (D).

**Figure S5.**
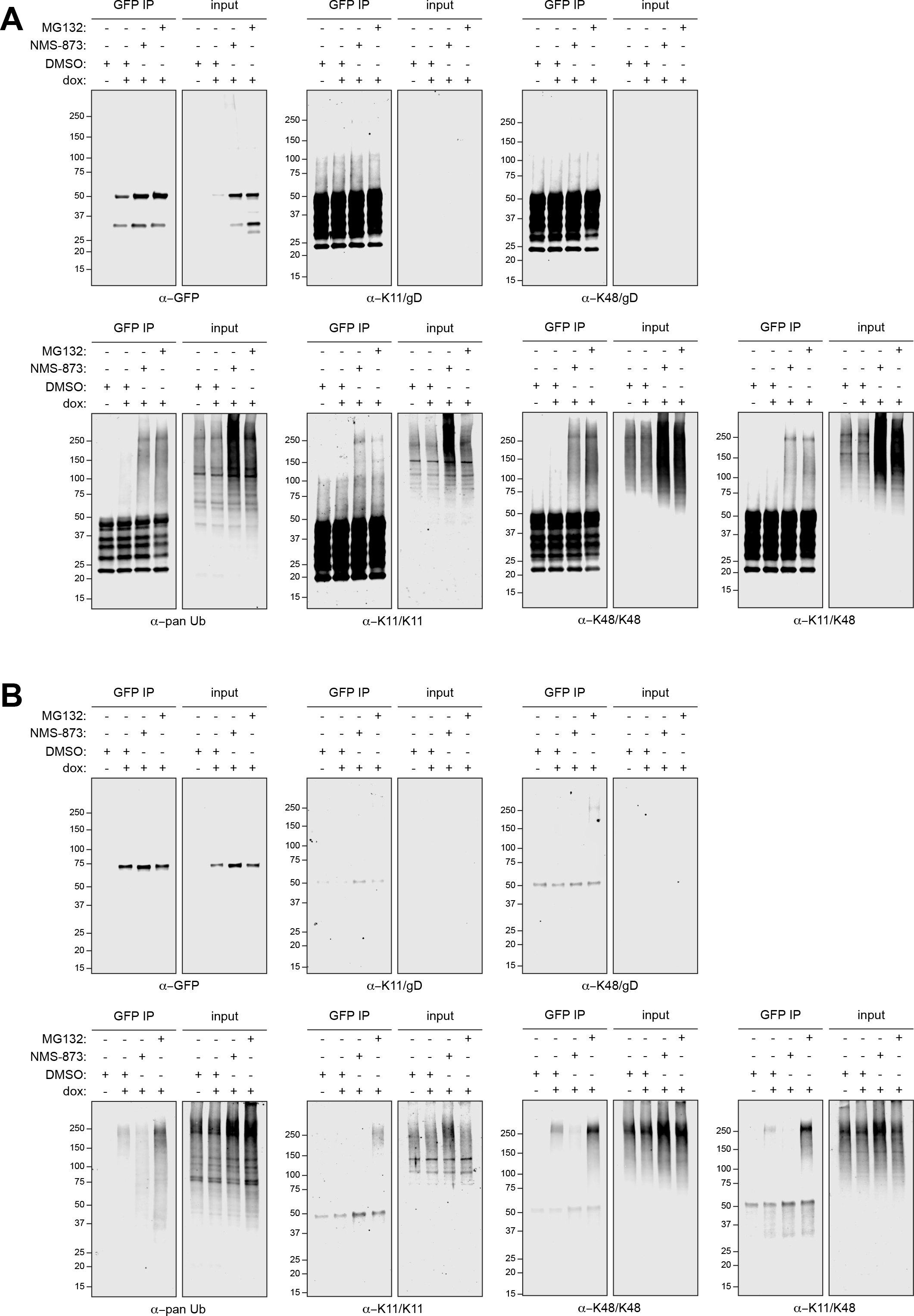
Detection of heterotypic Ub chains on ERAD-L and ERAD-M substrates, Related to Figure 5. **(A)** Full immunoblots and control immunoblots for Fig. 5D. **(B)** Full immoblots and control immunoblots for Fig. 5G.

**Figure S6.**
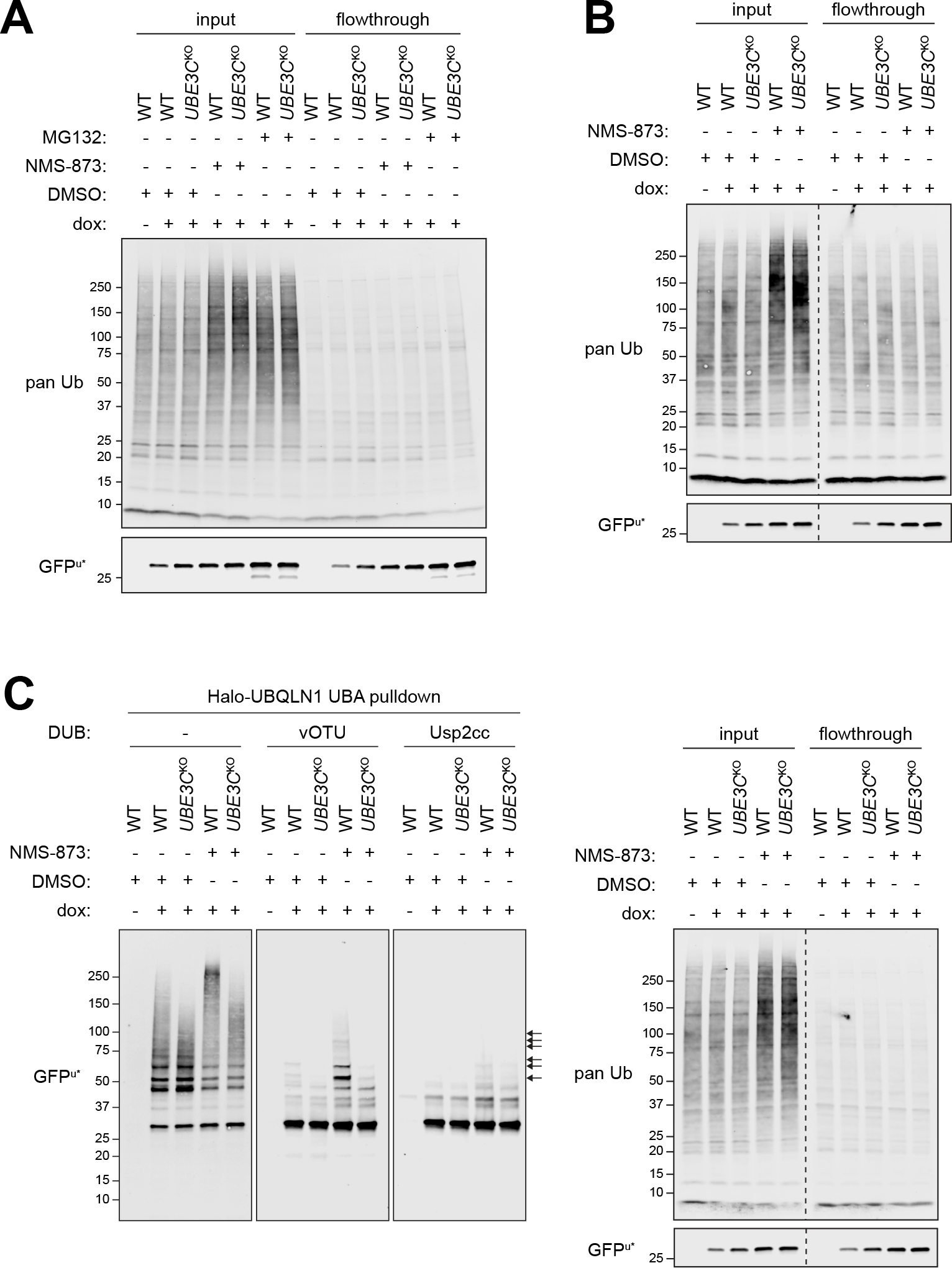
PolyUb-GFPu* is resistant to deubiquitylation by vOTU, Related to Figure 6. **(A)** Immunoblot analysis of cell lysate inputs and Halo-UBQLN1 UBA flowthroughs from Fig. 5B. **(B)** Immunoblot analysis of cell lysate inputs and Halo-TRABID NZF1 flowthroughs from Fig. 5E. Input and flowthrough samples were immunoblotted on the same membrane and are displayed with the same exposure time. Dashed line indicates removal of irrelevant gel lanes. **(C)** PolyUb conjugates were affinity captured from cell lysates using immobilized Halo-UBQLN1 UBA and the presence of K29 Ub linkages on GFP^u*^ was assessed by incubating with the catalytic domain of vOTU or Usp2cc as in Fig. 5H. Left: Ubiquitylated species were separated by SDS-PAGE and GFP^u*^ was visualized by immunoblotting with an anti-GFP antibody. Right: Immunoblot analysis of cell lysates before and after incubation with Halo-UBQLN1 UBA. Input and flowthrough samples were immunoblotted on the same membrane and are displayed with the same exposure. The dashed line indicates removal of irrelevant lanes.

**Table S1. Full casTLE results for each genome screen.**

**Table S2. Count files for each screen.**

**Table S3. Genes included in Figs. 2A-B, by category**

**Table S4. Ub peptides detected in INSIG1-GFP and A1AT^NHK^-GFP LC-MS/MS analysis**

**Table S5. sgRNA sequences used in study**

## Acknowledgments

We thank Amy Li and Gaelen Hess (Stanford University) for providing reagents and protocols, Alban Ordureau and Wade Harper (Harvard Medical School) for providing the Halo-TUBE recombinant protein, Marissa Matsumoto (Genentech) and Diane Haakonsen (University of California-Berkeley) for technical guidance for use of the Ub linkage and topology-specific antibodies, and Nicholas Pataki for help with R and Python packages. Sorting protocols were developed with help from Cathy Crumpton at the Stanford Shared FACS Facility (SSFF). Cell sorting and flow cytometry analysis was performed using instruments in the SSFF obtained using NIH S10 Shared Instrument Grants S10RR025518-01 and S10RR027431-01. This work was supported by a grant from the National Institute of General Medical Sciences (R01GM074874) and a generous gift from the Becker Family Foundation to RRK. DEL was supported by postdoctoral fellowships from the NIH (F32GM113370) and the Alpha-1 foundation. CW was supported by postdoctoral fellowships from the NIH ( F32GM113378) and Cystic Fibrosis Research, Inc (CFRI).

## Author Contributions

Conceptualization, R.R.K., D.E.L., M.C.B.; Methodology, D.E.L.; Formal Analysis, D.E.L., D.W.M., and L.Z.; Investigation, D.E.L., L.Z., and C.P.W. Writing-Original Draft, D.E.L. and R.R.K., Writing-Review & Editing, D.E.L., R.R.K., and D.W.M.; Visualization-D.E.L. and R.R.K.; Supervision-R.R.K., M.C.B., and J.E.E., Funding Acquisition-R.R.K. and D.E.L.

## Conflict of interest

The authors declare no conflicts of interest.

